# Effective computations for hippocampal place cell phenomena in sparse untrained random networks

**DOI:** 10.1101/2025.10.04.680487

**Authors:** José R. Hurtado, SueYeon Chung, André A. Fenton

## Abstract

The mammalian brain processes experience-dependent spatial information through poorly-understood network mechanisms thought to depend on particular network connectivity patterns and activity-dependent synaptic plasticity. However, dedicated input connections that learn to shape information about place cannot easily explain many rodent hippocampal place cell phenomena. For example, representational drift notwithstanding, the discharge of each place cell maps to specific locations in a fixed environment, but the discharge of most cells remap to distinct independent locations across environments, despite the fact that most sub-second cofiring relationships amongst hippocampal neuron pairs persist across environments. Whereas some models of hippocampal spatial information processing rely on the dedicated input connections of a for-purpose connectome (ignoring remapping, representational drift, and maintained cofiring), other models use synaptic plasticity implemented by learning rules to alter random input connections, but struggle with either limited capacity, representational drift, and/or biological implausibility. Here, using a randomly tuned network with feedback inhibition, we examine whether the assumptions of a specific connectome and learning-implemented synaptic plasticity are necessary for diverse place cell phenomena. We find that the random network with non-plastic connections accounts for positional tuning, single place fields in small spaces and multiple place fields in large spaces, mixed selectivity, and remapping, amongst other place cell phenomena. This requires excitatory activity to be sparse and organized across stimuli by divisive normalization. Enabling synaptic plasticity only at the network connections (not at network inputs) accounts for additional place cell phenomena including overdispersion, representational drift, and memory tagging. We show by simulations and analytically that DivSparse, a random network with sparsifying inhibition can explain many features of place cell network activity, suggesting that simple biologically-plausible architectures can realize representations of spatial experience that are robust, flexible, and spontaneous.

## INTRODUCTION

The much lauded cognitive map theory asserts that through the location-specific discharge of hippocampal neurons called place cells, the hippocampus forms an abstract map of the environment that uses the past to predict the future for avoiding threats and finding rewards, and making associations between important events and the places they occur (O’Keefe and Nadel, 1978). Place cells are hippocampal principal cells that selectively discharge in place fields, which are cell-specific locations of an environment (O’Keefe and Dostrovsky, 1971; O’Keefe, 1976; Muller, 1996). Most place cells have a single place field in environments with a maximum linear distance less than one meter, but in larger environments place cells express multiple irregularly-spaced place fields that can be somewhat enlarged (Fenton et al., 2008; Park et al., 2011; Rich et al., 2014; Harland et al., 2021). Hippocampal place tuning appears to be stable across multiple exposures to a fixed environment but when recordings are prolonged, the tuning is observed to drift with experience (Ziv et al., 2013; Hayashi, 2019; de Snoo et al., 2023; Geva et al., 2023; Levy et al., 2023). Despite the focus on place cell discharge, it is also important that in typical experimental environments, 70-80 % of hippocampal principal cells are either silent or express noisy spatial tuning, and that the most excitable cells tend to discharge as place cells (Thompson and Best, 1989; Kubie and Muller, 1991; Epsztein et al., 2011; Ziv et al., 2013; Stefanini et al., 2020; Levy et al., 2023). CA1 pyramidal cell firing is controlled by dominating inhibition, escape from which permits characteristic place cell discharge (Buzsaki, 1984; Kubie et al., 1990; Losonczy et al., 2010; Royer et al., 2012; Grienberger et al., 2017; Valero et al., 2022). Indeed, the place fields of a single place cell only occupy 15-20% of the environment such that ∼80% of the time a place cell is silent, only rarely discharging outside of its place fields, whereas inhibitory cells discharge ∼10 times faster everywhere in an environment, many with unique multimodal spatial firing rate tuning (Kubie et al., 1990; Fenton et al., 2000b).

Notably, changes in positional firing modulation of inhibitory cells predict the pattern of changes in the principal cell population, and presynaptic place cell firing fields modulate the positional tuning of interneurons, providing further evidence of the strong E-I coupling (Fenton et al., 2000a; Marshall et al., 2002). Firing restricted to place fields is remarkable because place cells receive hundreds of inputs from active projection neurons at every location (Olypher et al., 2002; Fenton, 2024). So much so that CA1 cells generate a new and persistent place field anywhere an experimenter delivers a transient ∼40 mV depolarization (Bittner et al., 2015; Bittner et al., 2017). Remarkably, place cells and their place fields are randomly redistributed when environments are sufficiently different, a phenomenon called “remapping” in which the internally-organized neural activity largely maintains, although its registration to the environment changes (Levy et al., 2023; Fenton, 2024). Because of remapping, the hippocampal space code is context-specific. Place fields are also modulated by attention, which along with context specificity is thought to underlie the contextual and episodic memory associations for which hippocampus is crucial (Kentros et al., 2004; Kentros, 2006; Mizumori, 2006; Mizumori et al., 2007; Muzzio et al., 2009; Smith and Bulkin, 2014; Pettit et al., 2022). Remapping challenges standard assumptions about the stability of stimulus tuning in the brain, as observed in sensory systems. Place cells can either remap (Markus et al., 1995) or fail to remap when tasks change in a fixed environment (Lenck-Santini et al., 2002; Lenck-Santini et al., 2005; van Dijk and Fenton, 2018). Another challenge to models is overdispersion, the fundamental unreliability of even excellent place tuning (Fenton and Muller, 1998; Olypher et al., 2002; Jackson and Redish, 2007; Fenton et al., 2010; Hok et al., 2013). Overdispersion is observed because hippocampal discharge is non-stationary. Place coding is multistable, organized according to the subject’s internal choice of spatial reference frame, a choice that can purposefully switch from one moment to the next in a manner that resembles attention and cognitive control (Gothard et al., 1996a; Gothard et al., 1996b; Fenton et al., 2010; Kelemen and Fenton, 2016; van Dijk and Fenton, 2018; Chung et al., 2021; Pettit et al., 2022). On the behavioral time scales of seconds and faster, position-related hippocampal discharge is multistable and overdispersed, although it is classically characterized by the minutes-scale phenomena of sparseness and spatial tuning measured by place fields that drift and remap demonstrating context-specific multistability. This collection of non-stationary features is challenging for models of place cell function that rely on dedicated for-purpose connections.

Most hypotheses explain place cells and their properties as learning-related changes in network connections, grounded in observations of activity-dependent synaptic plasticity in hippocampus (Bliss and Gardner-Medwin, 1973; Bliss and Collingridge, 1993; Mehta et al., 1997). As such, models assume that hippocampus learns a context-specific spatial map through experience-dependent adjustments of synaptic connections determined by context-specific patterns of coactivation during learning (Muller et al., 1996; Dobrunz and Stevens, 1999; Whittington et al., 2020). Such models encode connection strength between a pair of cells as the inverse distance between the two place field centers (Isaac et al., 2009), which creates place cell attractor network dynamics, as well as other experience-dependent phenomena (Blum and Abbott, 1996; Tsodyks et al., 1996; Samsonovich and McNaughton, 1997; Tsodyks, 1999; Mehta et al., 2000). Other place cell models rely on specially structured inputs with either specially structured input connections or activity-dependent learning rules that structure the input connections to generate geometric patterns (Zipser, 1985; Hartley et al., 2000; Lever et al., 2002; Solstad et al., 2006; Barry and Burgess, 2007). Other models begin with arbitrary connections and use activity-dependent learning rules to shape connections and learn a particular geometric map using either simple or complex multilayered networks (Muller et al., 1996; Whittington et al., 2020). Reinforcement learning is used to modify network connections in successor representations of networks designed to predict the likelihood of future network states given the current network state (Dayan, 1993; Gershman et al., 2012). Successor representations generate location-specific activity and other place cell phenomena when network states represent future locations, especially given basis inputs from elements with the spatial tuning of grid and boundary-vector cells (Stachenfeld et al., 2017; de Cothi and Barry, 2020). Thus a variety of network models have been proposed to generate place cell phenomena but all rely on structured inputs and/or connectivity, or synaptic plasticity to learn the connectivity, with varying degrees of biological plausibility.

The present work examines the assumptions of specialized functional connectivity and the necessity of synaptic plasticity for generating place cell phenomena by investigating randomly-connected networks with no synaptic plasticity. While hippocampal connectivity is not nearly as structured as neocortex, it is also not random (reviewed by Bernard and Wheal, 1994). Our contention is not that hippocampus is a randomly-connected network nor does it lack synaptic plasticity, rather this work examines the necessity of widely-held assumptions, especially in interpreting experimental work that these network features are essential. For example, it remains surprising that lesion of the medial entorhinal cortex (MEC) input to hippocampus does not alter CA1 place cell firing, although exciting the MEC layer 2 input causes remapping (Brun et al., 2002; Kanter et al., 2017; Robinson et al., 2017; Schlesiger et al., 2018). These inputs include grid cells, head-direction, border and non-spatial cells (Zhang et al., 2013) and CA1 activity appears to be more sensitive to direct MEC layer 3 inputs (Brun et al., 2001). The specialized and structured input from MEC is not essential for place cell firing, what about synaptic plasticity? Despite massive evidence of synaptic plasticity in the hippocampus, and evidence that maintained synaptic strength is essential for place field stability, a crucial role for synaptic plasticity in generating place cells is uncertain (McHugh et al., 1996; Rotenberg et al., 1996; Tonegawa et al., 1996; Wilson and Tonegawa, 1997; Cho et al., 1998; Kentros et al., 1998; Rotenberg et al., 2000; Nakazawa et al., 2003; Nakazawa et al., 2004; McHugh et al., 2007; Barry et al., 2012; Talbot et al., 2018), especially in light of so-called “engram cell” activation experiments that suggest synaptic plasticity is not required for memory storage or activation (Ryan et al., 2015; Pignatelli et al., 2019). Neurons that are genetically tagged by an excitatory opsin during memory expression have been called engram cells if their subsequent optogenetic activation is sufficient to express the memory (Ramirez et al., 2013; Tonegawa et al., 2015) because it is assumed that the cells are or at least contain the experience-dependent change that stores memory and defines the concept of an engram (Semon, 1921; Josselyn and Tonegawa, 2020).

We previously studied a randomly-connected neural network model with spike-timing dependent plasticity (STDP) rules that updated synaptic weights according to trajectories around a 1D ring environment, which generated positionally-tuned spiking activity of single network units with place cell properties (Levy et al., 2023). Neural dynamics in the spiking model were driven by feedforward inputs and feedback excitation (E) and inhibition (I) configured as a Leaky Integrate and Fire (LIF) network. We noted that inhibitory (EI and IE) plasticity played a crucial role in generating positional tuning, whereas EE plasticity did not. In fact, the plasticity itself is not required if feedback inhibition was fixed to silence all but a sparse and selective subset of cells (Fenton et al., 2023). These results suggested that network sparsity is essential for positional tuning, which is consistent with computational findings that inspire the present investigation (Pehlevan and Sompolinsky, 2014). While that work did not examine place cell properties, we were encouraged because it did generate sparse and abstract stimulus-selective network responses, with greater single unit selectivity in low-rate units (Pehlevan and Sompolinsky, 2014). Furthermore, theoretical work in odor-identity coding by the insect olfactory system suggests that sparse codes organized by random input connections are sufficient to generate high dimensional odor “tags” for downstream readouts (Dasgupta et al., 2017), and a similar model has been proposed to exist in the rodent olfactory system and the “caching barcodes” in the chickadee hippocampus (DasGupta and Waddell, 2008; Gill et al., 2020; Chettih et al., 2024).

This computational mechanism, called “locality-sensitive hashing,” was recently leveraged to model associative memories between the hippocampus and MEC networks while integrating external sensory cues, producing robust benefits in terms of storage capacity and the completeness of recalled representations (Chandra et al., 2025). The locality sensitive hashing function in the temporal domain of memory representations relies on fixed random input connections from the MEC to HPC networks. Moreover, to explain time-averaged place cell firing fields, inputs to the hippocampal CA1 region have been effectively modeled as random Gaussian processes followed by simple post-processing steps, where plasticity may only play minor additional roles (Mainali et al., 2025).

In the current work, we investigate the expressivity of random connections in an LIF network model, motivated by results from the model showing that random input connections and feedback inhibition are sufficient to induce place representations that are capable of remapping (Levy et al., 2023). Using the same EI network model, the present work aims to determine how many diverse hippocampal phenomena can be explained in biologically realistic spiking and rate-based networks that do not require learning, plasticity, or global objective functions that constrain the connectivity. This can help to segregate which phenomena of the hippocampal network may be a result of its learning functions and which may be pre-existing substrates for learning, despite the same molecular mechanisms contributing to both (O’Reilly et al., 2018; Chung et al., 2021). In our previous work, once the network connectivity weights were set through STDP learning rules, they no longer needed to change to generate novel place cell responses when the inputs remapped (Levy et al., 2023), in contrast to many but not all models of place cell location-selectivity that rely on learned or bespoke EE connections (Blum and Abbott, 1996; Muller et al., 1996; Barry et al., 2006; Solstad et al., 2006) but consistent with the computational findings (Pehlevan and Sompolinsky, 2014) and the empirical effects of manipulating synaptic plasticity and hippocampal inputs reviewed above. Using both LIF network simulations and an equivalent analytical “DivSparse” model, the present work demonstrates that the network generates all the place cell phenomena reviewed above, so long as the inhibition is optimally set to generate sparse network activity. By enabling local synaptic plasticity to act on the pre-existing network we demonstrate how the network can enable place and threat avoidance learning and memory recall, as well as alternating reference-frame specific spatial representations according to attention-like cognitive control signals (Table 1). It is especially noteworthy that the work demonstrates it is possible to generate phenomena that empirical studies use to identify memory, sometimes interpreting these phenomena as evidence that experience causes enduring network changes identified as “engrams,” and “engram cells,” without in fact any such changes needing to occur.

**Table 1:**
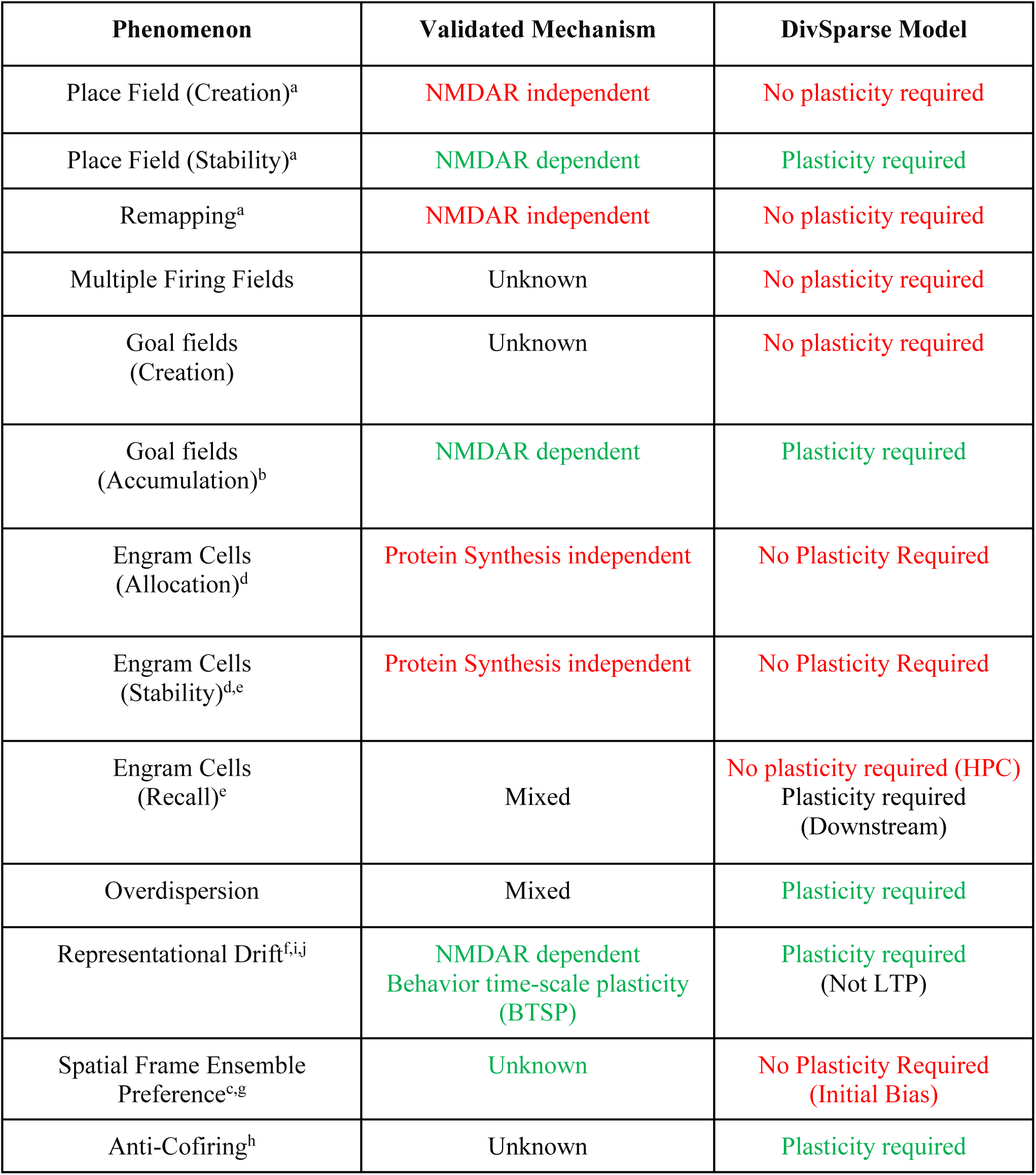
A list of experimental phenomena in HPC and their associated hypotheses. Green and red text indicate that plasticity is or is not required, respectively. References: ^a^Kentros et al. 1998, ^b^Dupret et al. 2010, ^c^Kelemen and Fenton 2010, ^d^Ryan et al. 2015, ^e^Pignatelli et al 2019, ^f^Hayashi et al 2019, ^g^Chung et al 2021, ^h^Levy et al 2023, ^i^de Snoo et al 2023. ^j^Maddar et al 2023.

## RESULTS

### Place field formation and remapping in a random untrained network with feedback inhibition

We begin by investigating whether an EI network with feedforward all-to-all random input connections (Fig. 1A,B) can generate place cell-like (position-tuned) responses without adjusting input or network connections. In contrast to conventional EI-balanced networks, the network is not constrained to be explicitly EI-balanced (i.e. symmetric inputs to E and I populations with reciprocally balanced weights). Instead, all EI and IE weights are preset to the same value for simplicity (Fig. 1A-I) or randomly assigned from a scaled uniform distribution (Fig. S1). Recurrent EE and II weights were set to zero to simplify the analysis, motivated by previous work showing that these connections are not required for place-specific responses (Levy et al., 2023). Input spikes are modeled as inhomogeneous Poisson spikes that respond to the current trajectory using a deterministic Gaussian mixture model, where each input contains on average six overlapping Gaussian distributions that are randomly positioned. Since the inputs derive from many random positions, and since the feedforward connections are all-to-all onto E neurons, the information content of the resulting network outputs should lack single-cell selectivity in the absence of feedback inhibition. We therefore performed simulations across a range of preset EI and IE connections, showing an optimal range of feedback inhibition for generating activity-dependent positional tuning (Fig. 1C,G, Fig. S1-S3). Although formally, the network is not EI balanced, interesting phenomena emerge that resemble explicitly EI balanced networks (Fig. 1A-D). The single-cell place selectivity is predicted by a three-stage computation: 1) a linear stage multiplying the binary input peak location matrix of the input tuning by the matrix of random feedforward input connections, 2) normalization across network E neurons (Fig 1B, left) and 3) thresholding weak activations to zero. Sorting neurons according to their strongest input location following normalization reveals the underlying positional tuning that emerges in the LIF membrane potential dynamics (Fig. 1B right). The nonlinear components of this transformation arise from the leaky membrane dynamics, the threshold-spiking mechanism, and the feedback inhibition. Importantly, spontaneous place selectivity arises once the network is sufficiently sparse (Fig. 1C,D). The output neurons that are most sensitive to the current input pattern cause inhibitory feedback to decrease the likelihood of other neurons spiking at that location, thus, when inhibitory strength sets the network slightly above critical sparsity (Fig. 1D), positional tuning emerges spontaneously and therefore remaps naturally when the environmental inputs change (Fig. 1E). If there are enough excitatory neurons in the network, the optimal inhibitory strength approaches critical sparsity (Fig. S2), suggesting that near the phase transition of critical sparsity, small inhibitory changes have outsized effects on selectivity and sparsity in the network. Such an inhibitory regime is important for neuronal avalanches, as well as other spontaneous neural processes (Beggs and Plenz, 2003; Buzsaki et al., 2007; Sanzeni et al., 2020; Sanzeni et al., 2023).

**Figure 1:**
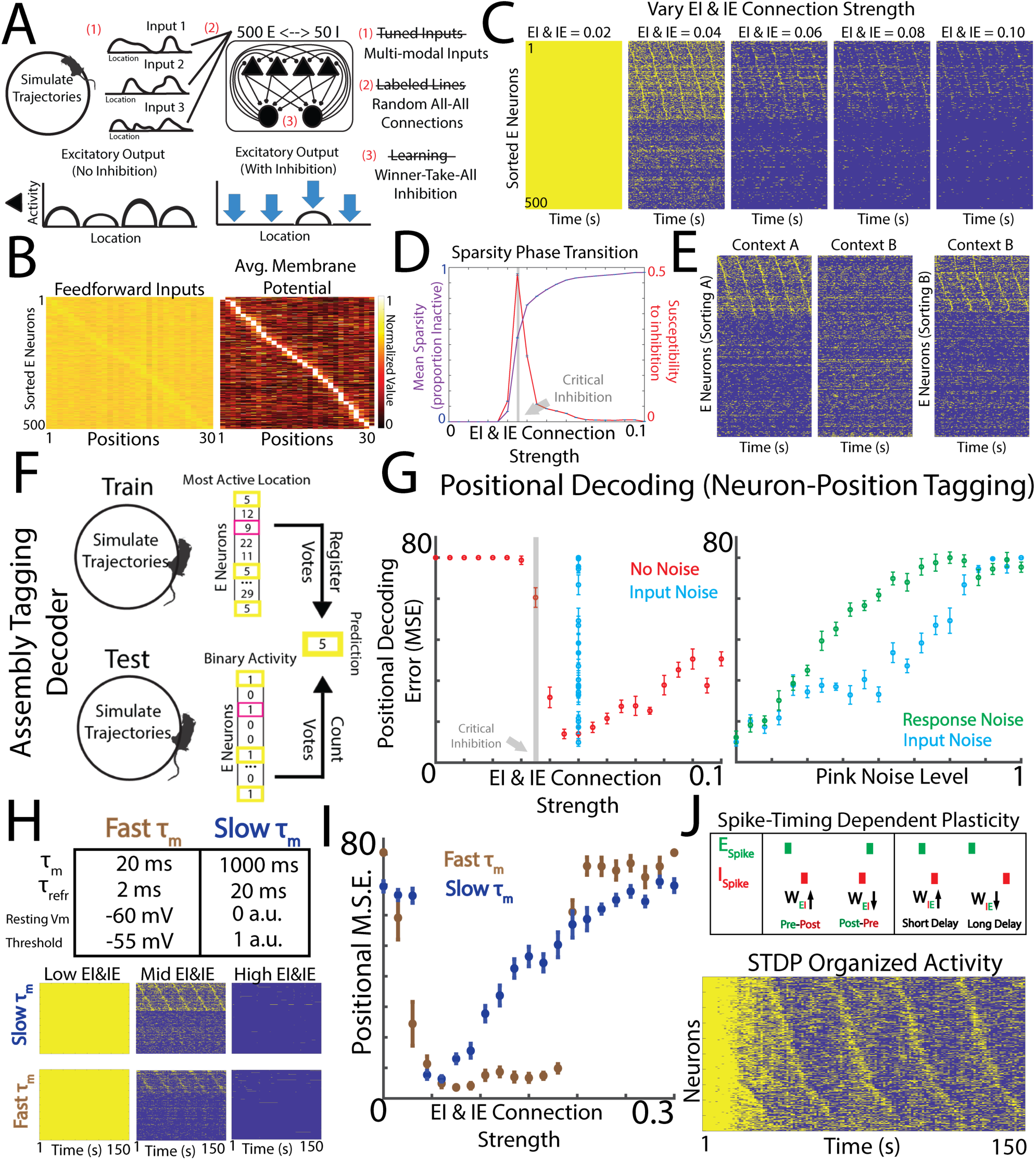
Positional tuning and remapping in an untrained network using inputs with random all-to-all connections. A) Schematic of EI network architecture and underlying assumptions. Top Left: Inputs are sampled as random multimodal positional filters that draw spikes from simulated trajectories around a circle. Top middle: Random all-to-all input connections mix the input activities across the network E neurons while I neurons provide feedback inhibition. Bottom Left: The response of a network E neuron without inhibition is schematized as random bumps of activity in multiple locations. Bottom Middle: the response of the same E neuron with feedback inhibition shown as a single bump of activity and the emergent positional tuning in the E neuron. Right: Assumptions of the model and deviations from standard assumptions. B) Left: Example linear drive onto the E cells from the product of the input filters and random connections, peak-normalized at each position across E cells, sorted according to the peak drive location. Right: The LIF with inhibition network response selects location-specific peaks, illustrated by the trial-averaged membrane potential at each location after min-max normalizing. C) Spiking activity of simulations as a result of increasing inhibitory connection strength. D) Susceptibility of sparseness to changes in inhibition is shown as the derivative of sparseness with respect to changes in inhibitory strength. The peak in susceptibility indicates a phase transition in Sparsity at inhibition values of ∼0.03. E) Defining new contexts by randomly redrawing the input filters, the network remaps, changing the location preference of the E neurons in the network. Shown are output spikes in context A and context B, and only when the sorting matches the context is the appropriate map revealed. F) Assembly tagging decoder used to enforce a metric of sparse activity patterns. Trial-averaged positional tuning is determined in a previous training trial, then during a testing simulation, each time the neuron is active is a vote for its preferred location. The location with the most votes is the decoded position, therefore activity must be sparse and based on positionally tuned cell assemblies. Such a decoder is meant to capture activity-dependent codes related to memory in the hippocampus, which uses a remarkably sparse code. G) Effect of increasing inhibition (left) and increasing input and response noise (left and right) in the assembly tagging decoder. Note the robustness of the EI network to input noise up to the noise level where input and response noise are equal. H) EI model parameters can be altered while preserving the effect of untrained positional tuning (see activities for the network with biological parameters). I) Decoding error for biological and nonbiological EI model parameters, note the optimal inhibitory strength that minimizes decoding error in the untrained network. J) Top: Spike-Timing dependent plasticity rule (STDP) for EI and IE synapses in the LIF model. Bottom: LIF Network simulation using STDP to generate positional selectivity in an unsupervised manner. Neuron sorting was computed prior to the simulation using stimulus-specific sorting of the expected feedforward linear responses across E neurons.

Sparse location-specific discharge of hippocampal neurons identifies active subpopulations that can represent the current position in an environment (Brown et al., 1998). We therefore analyzed positional tuning in this LIF network using an assembly tagging decoder (Fig. 1F,G) that identifies each neuron’s preferred location during the training phase. Each neuron’s preferred location is a vote casted whenever the neuron is active during the testing. At each test time t, the location with the most votes across neurons is the decoded location. Since positional information is distributed across the activations regardless of sparsity, a linear decoder is sufficient to reasonably decode location even without inhibition (Fig. S3). However, the assembly tagging decoder penalizes these activations, since every neuron is active in every location. Instead, the assembly tagging decoder emphasizes sparse subsets of cells that can be associated with experiences in a selective code that is sparse. Furthermore, in the sparse regime, the assembly tagging decoder outperforms linear regression with or without regularization (L1 or L2) (Fig. S3). Although the real hippocampal code surely breaks the assumption of a dedicated code, particularly in large environments that generate multiple place fields per neuron, most experiments that we aim to reproduce in this work were performed in small environments under the assumption of a dedicated code that could facilitate assembly tagging (Fenton et al., 2008; Park et al., 2011; Rich et al., 2014; Harland et al., 2021). Under these assumptions, an assembly tagging decoder is sufficient to quantify the network performance, which is our present goal.

Importantly, the assembly tagging decoder shows that the LIF model compresses the activity of a highly distributed, globally active input signal onto far fewer and sparser output cells that are more selective than the inputs (Fig. S3C-F). This results in more information per spike due to the combined effects of sparsification and compression, lending measurable benefits to neuronal metabolism and energy expenditure (Laughlin et al., 1998). In fact, decoding via assembly tagging on the LIF activity outperforms direct decoding of the inputs using either decoder (Fig. S3F). Therefore, a readout mechanism adapted for sparse codes (like an assembly tagging decoder) would be enhanced by the same EI sparsification mechanisms that can generate spontaneous remapping and selectivity from highly distributed inputs (Fig. S3E,F).

Furthermore, the network is also amenable to unsupervised tagging mechanisms more explicitly tied to memory encoding and reactivation experiments (Fig. S5, S6). During these simulations, activity dependent tagging of neurons occurred at special locations marked by using transient increases in input activity as a training signal (Fig. S5C). A subsequent simulation without the training signal can test whether a downstream readout can associate the tagged assembly with an approach/avoid response that was selective of the special location (Fig. S5D). This readout mechanism works without the need for labeled decoding, and it is more analogous to memory tagging and reactivation experiments with activity dependent immediate early genes (IEGs). (Fig. S5, S6). Moreover, contextual decoding was simultaneously performed by tagging assemblies according to primacy (first neurons to N spikes) within a given behavioral session (Fig. S5A, B). Primacy has been studied as a plausible way to readout temporal codes, particularly in olfaction (Wilson et al., 2017), and serves the role of a contextual readout mechanism in our model to complement the positional readouts. In this way, the pairing of the EI network and downstream reader is capable of behavior that resembles active place avoidance and contextual fear conditioning (Fig. S6) from the same activations, all in the absence of plasticity in the encoding layer. It remains to be seen how plasticity mechanisms can be leveraged to enhance memory encoding for multiclass tasks, as well as the constraints on memory capacity. These findings show how an untrained network with random input connections and feedback inhibition can generate efficient positional tuning and remapping using binary activations of E subpopulations, once levels of feedback inhibition are adequate. These results suggest that the presence of an unlearned and reliable spatial map of any arbitrary context can allow associative learning to propagate effects of memory-associated units with the features of what are described as “engram cells,” but without requiring any alterations to the intrinsic mapping function (i.e. the random input connections) but at the same time permitting feedback inhibition to regulate the nonlinear transformations of the network. This shows how the term “engram” to describe tagged cells can contradict the fundamental concept of “engram” as an enduring change (Semon, 1921).

It is likely that the LIF model has several parameters unrelated to the HPC-like phenomena of interest, therefore we implemented versions of the LIF model with very distinct network parameters. Across distinct sets of possible model parameters, which can be biologically plausible or implausible, the phenomenon of optimal inhibition also emerges for positional tuning (Fig. 1H,I). This suggests that the specifics of the temporal dynamics of the model are less important than their ultimate effect on the network’s inhibition-mediated selectivity. Since average inhibitory effects are more important than specific inhibitory connections (Fig. S1), we next investigated whether unsupervised plasticity rules can self-organize the inhibition to reproduce the spatial maps expected from the optimal inhibition. We therefore allowed spike-timing dependent plasticity (STDP) rules to alter the EI and IE connections, making the network self-regulating (Fig. 1J, see Methods). Even after initializing the EI and IE weights at an unstable state (arbitrarily low). The STDP rules modify the weights to bring the network to a sufficiently balanced output, revealing the intrinsic positional tuning (and neuronal sequences) computed prior to the simulation directly from the inputs and random connections (Fig. 1J). Given the importance of synaptic plasticity for hippocampal function, it is important to be clear, the STDP in the LIF model is inhibitory, not excitatory, and allows for self-organized representations that do not require hand-tuned inhibition (Fig. 1J), making the network self-organizing, and its representations robust to arbitrary initializations. Next, we examine which features of the LIF model can be abstracted into a simpler model architecture that is more analytically tractable while reproducing the same relevant phenomena from the biologically plausible and more complex spiking model.

### Normalization and thresholding of random tuning in output neurons explains stimulus tuning features of LIF model

The relevant hippocampal phenomena of place tuning and remapping was reproduced in the LIF model despite the model being in very distinct parameter regimes or using unsupervised plasticity mechanisms. Therefore, we next investigated whether the temporal dynamics and specifics of the inhibitory connections can be abstracted and simplified in an interpretable model. We hypothesized that context-specific positional tuning is the result of three simple sufficiency conditions: 1) broad selectivity of input neurons, 2) moment-to-moment normalization of outputs, and 3) a threshold that silences all but the most selective subpopulation at any moment. Using the same input connectivity and stimulus encoding as the LIF model (Fig. 2A), we abstracted away most features of the EI network, such as the input dynamics, membrane potentials, membrane time constants, feedback inhibition, and refractory periods. The simplified model called Divisive Sparsification (DivSparse) consists of max-normalizing feedforward inputs across the pool of output E neurons (a form of divisive normalization) and setting a threshold at every stimulus (Fig. 2B).

**Figure 2:**
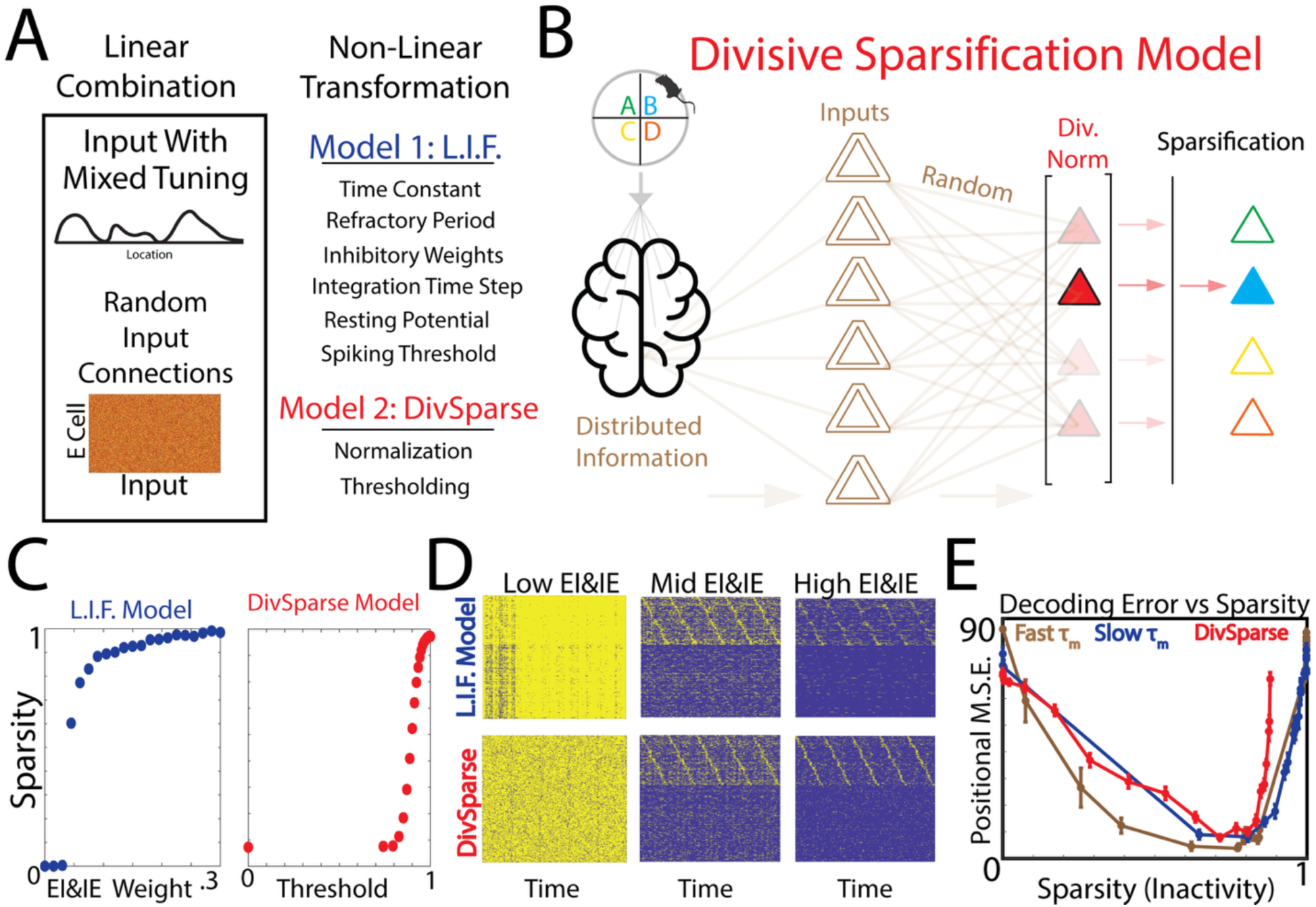
Simplified and more interpretable DivSparse network model reproduces features of EI network. A) The tuning curves for the DivSparse are generated from (Left) a linear combination of input filters and random connections and (right) normalization across neurons at each stimulus and a sparsifying threshold. B) Schematic of the DivSparse network. Non-zero values after thresholding can be used as rates in a rate model, or they can define an inhomogeneous Poisson spiking model as done below. C) Comparison of EI model sparsity and the sparsity of the simple model as a function of inhibition weights and threshold values, respectively. Increasing inhibitory strength in the EI model is analogous to increasing the threshold in the simple model. D) Spiking activity in the two models. E) Decoding performance as a function of sparsity. Note there is a shared sparsity that achieves optimal decoding (∼80% Inactive Units).

To compare the two models, we generated inhomogeneous Poisson spikes with the sampled spike rates given by the DivSparse activations. The key difference between the inhomogeneous Poisson spikes in the LIF model and the DivSparse model is that the LIF model uses the inhomogeneous Poisson process to generate <u>input</u> spikes which drive the LIF network spikes. However, in the DivSparse model, the Poisson process is directly generating the network <u>outputs</u>, using the non-zero entries of the DivSparse activations as centers and peaks of a Gaussian mixture model (See Methods). Increasing the strength of inhibitory connections in the LIF model is analogous to increasing the threshold in the DivSparse model (Fig. 2C), as both increase the sparsity (proportion of inactive units) in the output activity. Using the same input tuning and input connections, the simplified model reproduces the spatial tuning of the LIF model (Fig. 2D). Moreover, the analogy becomes clear when comparing the positional decoding of both models as a function of the network sparsity (Fig. 2E). The DivSparse and the two distinct LIF models result in a similar optimal sparsity that minimizes decoding error (∼80% inactive units). These findings suggest that network sparsity is the critical parameter of interest that is modulated by an effective thresholding of normalized activities. This arises in the LIF simulations by a combined influence of the feedback inhibition, the leaky membrane potential, and the spiking nonlinearity while arising in the DivSparse model from divisive normalization and the sparsifying threshold. We next explore the multimodal capabilities of the DivSparse model, where inputs contain both spatial and nonspatial signals (landmarks or sensory) and often contain multiple competing spatial frames of reference.

### DivSparse model generates multimodal, multi-frame representations if divisive normalization occurs independently across modalities

We next examined whether the DivSparse model produces multimodal and multi-frame place representations as observed during rodent navigation (Fenton et al., 2010; Kelemen and Fenton, 2010). First, we simplified the spiking DivSparse model into a rate model by ignoring the Poisson spike generating step. Instead, the DivSparse activations directly encode the rates sampled at a given location. We then expanded the inputs of the model to include correlations along the signaling domain (position), such that inputs at nearby locations generate more similar patterns. Without divisive normalization, the sparsification step acts discontinuously across the stimuli, preventing selectivity across the stimulus domain (Fig. 3A). On the other hand, with divisive normalization, sparsification leads to selective activity regardless of the presence or absence of input signal correlations. Furthermore, different naturalistic stimuli contain distinct correlation structures across their signaling domain (i.e. adjacent positions). For instance, spatial inputs into the hippocampus are trivially correlated along the signaling dimension of positional signals. However, this is not necessarily the case for objects, textures, landmarks and other stimuli that lack extended spatial autocorrelations. To explore the effect of combining uncorrelated landmark signals with correlated spatial signals under conflicting spatial reference frames, these inputs were combined to generate multimodal, multi-frame representations that are typical of hippocampus and best observed when continuous rotation of the behavioral arena dissociates the rotating arena frame of reference from the stationary room frame of reference (Fig. 3B; Fenton et al., 1998; Fenton et al., 2010; Kelemen and Fenton, 2010, 2016). The positional signals were modeled as positionally correlated inputs, as would be expected for grid cells, head-direction cells, and other spatially modulated inputs to hippocampus (Fig. 3C).

**Figure 3:**
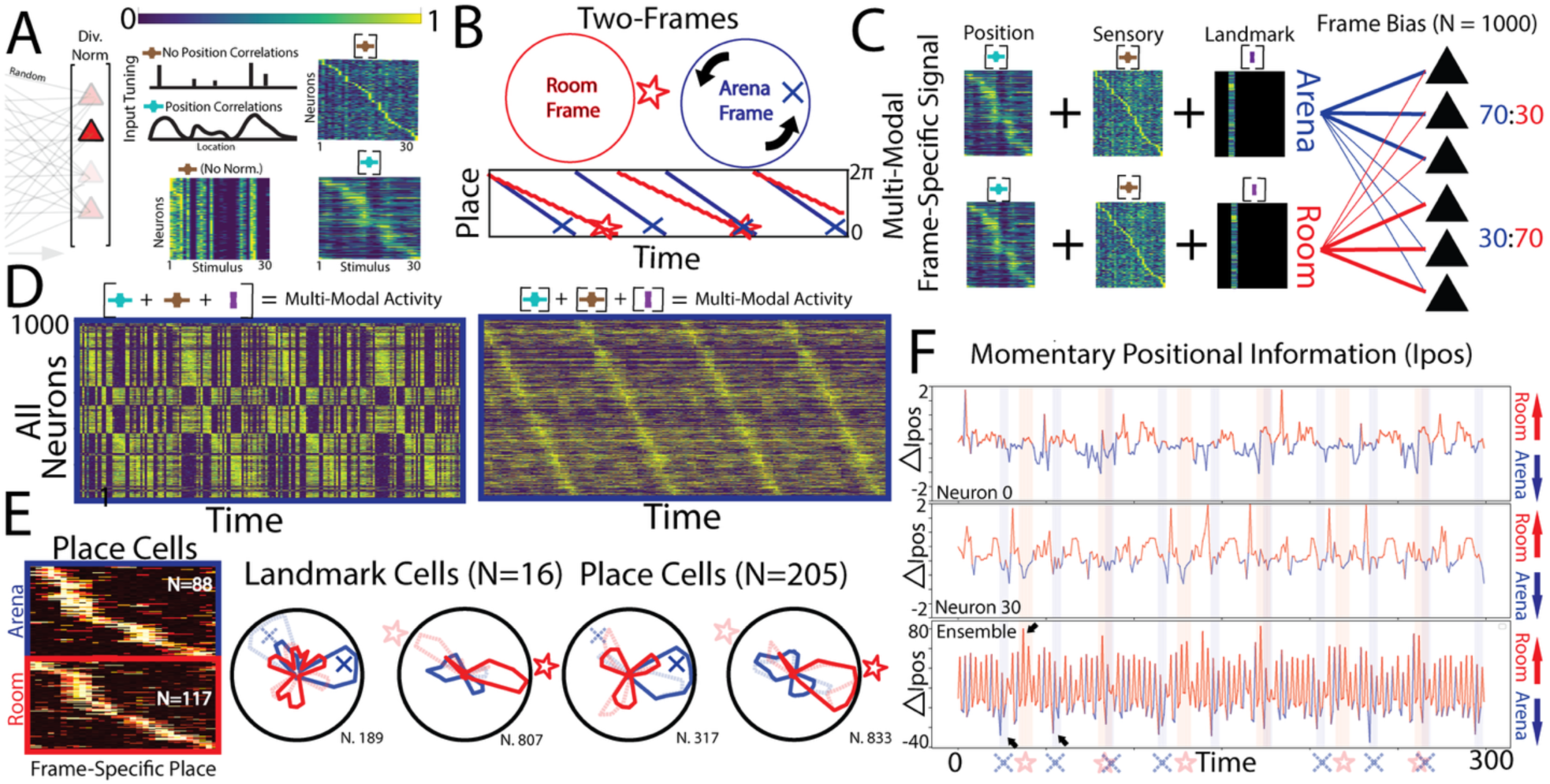
Divisive normalization and sparsification enables multimodal, multi-frame representations. A) The divisive normalization (Left), and the input correlations across positions (Top Middle), can be included or removed to study the effect on the model. Bottom Midde: The model without a divisive normalization step (only thresholding the feedforward inputs) fails to generate selective activity. Right: The DivSparse model without input correlations (Top) and with input correlations (Bottom). B) Simulated trajectories over two simultaneous spatial frames: a stationary room frame (red) and a frame fixed to a rotating arena (blue). Special landmarks are also placed at special locations in each frame to test for landmark anchoring. C) From these simulated trajectories, a multimodal frame-specific matrix is generated by combining randomly generated DivSparse outputs with different correlation structures (see (A)). The two distinct multimodal frame specific matrices are then assigned distinct frames (room vs arena) as samples are drawn from the columns of each respective frame according to the simulated trajectories (From B). A 70:30 Arena:Room input bias was enforced on all inputs onto half of the neurons and a 30:70 bias was enforced on the other half. D) Network activity across all neurons in the multi-frame simulation. The neurons are sorted by preferred Arena Frame location. Divisive normalization after (Left) versus before (Right) adding together the inputs produces different effects. Only divisive normalization before addition (Right) leads to smooth representations of the trajectories. E) Left: Place cells in both frames extracted from the multi-frame model in D. Right: example of landmark anchoring and place cell tuning curves. F) These representations dynamically toggle between Room and Arena frame preferring states at the single neuron level (top, middle). Bottom: The ensemble positional information also switches preferred frame dynamically, particularly when a frame-specific landmark is encountered in either frame (black arrows).

Background sensory signals were modeled as uncorrelated inputs placed in locations distributed across the 30 positions, representing any signals that may arrive due to sensory features of a given location while lacking broad correlations with nearby locations (Fig. 3C). Lastly, landmark objects were generated as a single stimulus placed over three adjacent locations, representing the extent of the landmark (Fig. 3C). To generate two dissociated spatial frames, two distinct versions of this setup were generated and coupled position-wise, one defining an arena frame and the other defining a room frame of position-sensory-landmark representations. To modulate the combination of DivSparse models, a bias was enforced such that half of the neurons received stronger arena frame inputs (70:30 bias) while the other half received stronger room frame inputs (Fig. 3C). Trajectories around a circle are then used to generate activities from each frame, which is then combined via addition prior to sparsification. The resulting model is a full multi-frame, thresholded matrix of activities. With the multimodal signals, there is a choice of applying divisive normalization before or after combining the different inputs, and the resulting model was examined separately with each normalization scheme (Fig. 3D). Rotation of the arena is modeled by a constant phase shift that shifts the position-position coupling between the room and arena frame while sampling activities from the trajectories. Surprisingly, divisive normalization before summation and sparsification is the only scheme that leads to smooth spatial representations capable of anchoring to both positions and landmarks in either frame using a special subset of cells (Fig. 3D, E). The time series of frame-specific positional information demonstrates that both individual neurons and the ensemble (as a coordinated population) transiently switch between representing positions in the room and arena frames (Fig. 3E,F). Moreover, the ensemble positional information transiently switches to the frame of whatever landmark was recently encountered. This reproduces experimental observations during two-frame active place avoidance tasks, driven by explicitly sensory inputs rather than network intention or cognitive control. In summary, the DivSparse model produces multimodal, multi-frame representations flexible to choices of input types, correlation structure, divisive normalization, and sparsity. These choices are especially relevant to understanding the requirements that allows or prevents a downstream readout or tagging mechanisms that may occur downstream of the hippocampus.

### Simplified 2D model of positional tuning with local plasticity reproduces several additional features of hippocampal place cell activity

The 1D DivSparse model was extended to two dimensions with some additional parameter choices. The stimulus domain was made 2-dimensional, resulting in 2D DivSparse activations. To smooth the activations in the network, a 2D Gaussian mixture model was wrapped over the non-zero activations (centered at each nonzero value with peaks defined by the nonzero value and using a randomly sampled Gaussian width) (Fig. 4 A,B). We next examine the features of the 2D DivSparse model, and how it compares to known features of hippocampal activity in 2D environments. Since the simplified 2D model generates firing fields from the random tuning and connections coming from the inputs, several features of the model, such as remapping (Fig. 4B) could be qualitatively compared to experimental recordings. In one set of simulations, the environment from which trajectories were sampled was enlarged, and in accordance with experimental findings, there are cells with single firing fields in one environment that generate enlarged, multiple, and aperiodic fields in the larger environment (Figs. 4B, S5, S8; Fenton et al., 2008; Park et al., 2011; Harland et al., 2021). This model result is straightforward. For any cell, every sampled unit of area contains a nonzero probability of generating a place field. Therefore, the larger the area, the larger the likelihood of generating fields (Fig. 4C). This reproduces a finding that was recently made statistically rigorous in the framework of Gaussian processes (Mainali et al., 2025). Next, we examine the expressivity of spatial representations when local plasticity is allowed to occur between recurrent EE connections, akin to the plasticity found in Ammon’s horn of the hippocampus. The local plasticity operates within a short time window in an activity dependent manner according to spike-timing-dependent plasticity rules, reinforcing connections between neurons that fire earlier onto those that fire later, while decreasing the reverse connections (Dobrunz and Stevens, 1999). We found that this local plasticity allows the firing fields of individual neurons to drift moment to moment, leading to overdispersed positional tuning while retaining the same trial average positional tuning (Figs. 4D-E, S8, S10). To explore the combination of local plasticity and synaptic potentiation, an “LTP Tagging” signal was included (Fig. 4F) to strengthen recurrent connections amongst neurons coactive during the brief crossing of some special location (defined by a behaviorally relevant signal like a shock or reward). The LTP across a subset of cells in the network led to a stabilization of the drifting phenomenon as the firing fields reorganize (Fig. 4G), while other fields formed near the tagged location, much like the so-called “goal cell” firing fields described in hippocampal experimental recordings (Hollup et al., 2001; Hok et al., 2007; Dupret et al., 2010).

**Figure 4:**
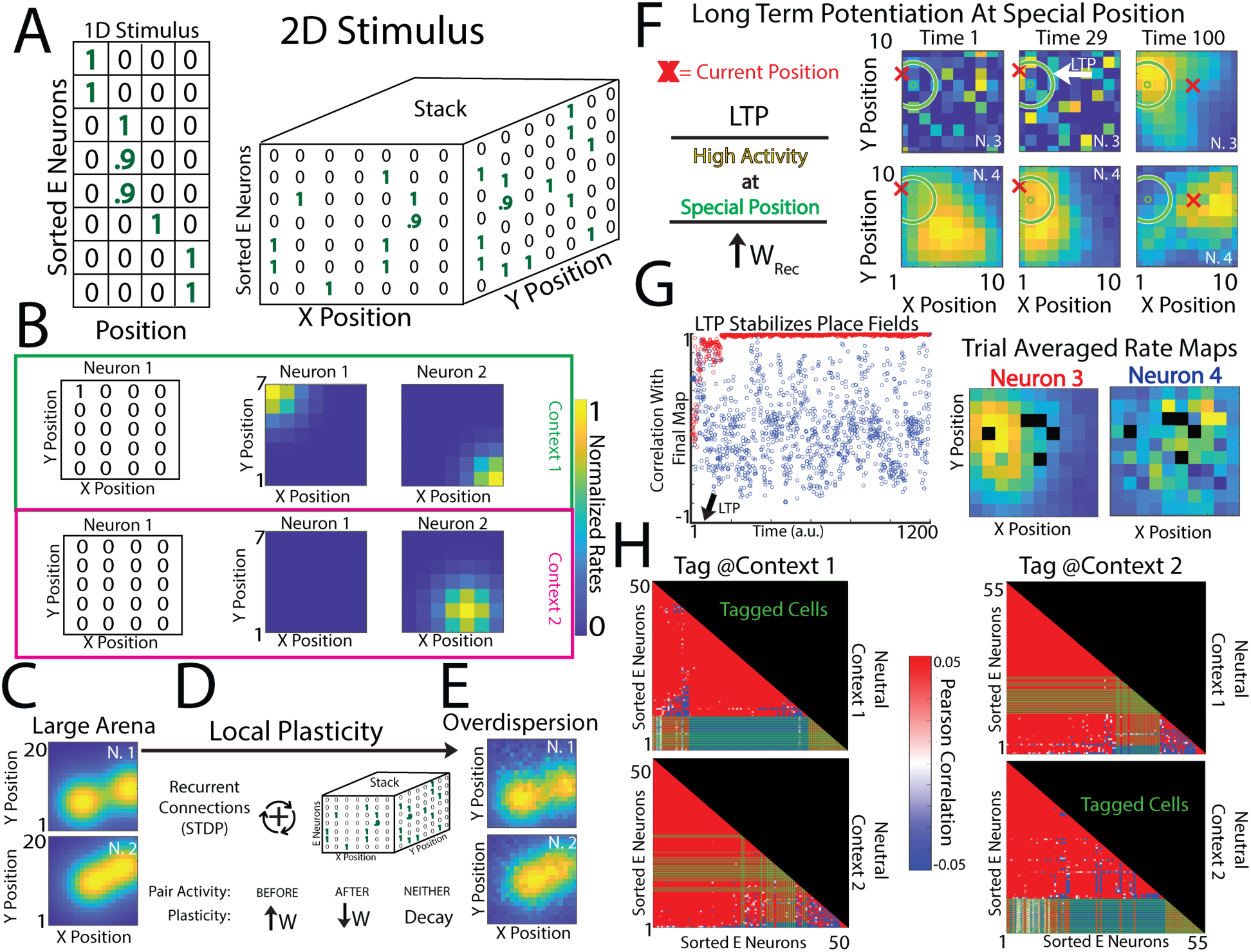
The DivSparse model can generate 2D place fields expressing idiosyncratic features of rodent hippocampal activity. A) 2D place fields were generated from the DivSparse model by only changing the 1D to 2D input filters. The 2D tuning curves of all the output neurons can be stacked into a 3D matrix as shown on the right. B) Example of trial-averaged simulated activity for two example neurons (thresholded input filters and Gaussian-smoothed firing fields shown) in two contexts, that show remapping. C) Multiple place fields emerge when the environment increases from 7 (red) to 20 unit squares on a side. D) Linearly combining neuron tuning curves using the recurrent weights and adding local plasticity to the recurrent connections (initially set to 0). An asymmetric spike-timing-dependent plasticity (STDP) rule set the weights such that weight w_ij_ increases when i fires earlier than j and decreases otherwise. This local plasticity causes several idiosyncratic phenomena (Fig. S8). E) Overdispersion of location-specific activity (compare to C) while preserving trial-average place field centers. F) Learning-induced place fields at special location: the activity of two neurons undergoing local plasticity drifts (compare distinct time points), one of which (N3, t=29) generates a new place field when activity (red “x”) is coincident at a special location (green circle), at which point the wij’s increase for all coactive neurons i and j. G) Left: N3’s Firing rate map stabilizes after LTP, while N4’s does not. Right: Trial-averaged firing rate maps of the two neurons from (F). Right: Final map stability - 2D Pearson correlation between a neuron’s firing rate map at time t and the last time point. The correlation increases and is maintained for the neuron with the LTP-stabilized place field. H) Pearson correlations of neurons during two neutral contexts (context 1 and context 2) after location-specific LTP has occurred in one of the contexts. Neurons are sorted according to anti-cofiring power (proportion of pairwise correlations < -0.05). Green lines indicate the tagged LTP-neurons. (See Fig. S9 for more examples).

Lastly, we compared the cofiring properties of the tagged and untagged cells in the DivSparse model with local plasticity and LTP tagging. Due to the positional bias inherited by a subpopulation of neurons coactive at the tagged location, these neurons develop anti-cofiring statistics which facilitate contextual discrimination (Figs. 4H, S9). This is similar to experimental results in freely-behaving mice (Fig. S8; Levy et al., 2023). Thus, the presence of spatial maps generated from common input connections of distinct strengths can allow for synaptic mechanisms that transform maps in ways that reproduce several peculiar phenomena of hippocampal place cells.

## DISCUSSION

### Summary and Limitations

The current findings provide a unifying model to understand the spatial processing of a toy network that reproduces many features of hippocampal place cell activity (Table 1). Importantly the framework does not adopt long-held assumptions of specific connectomes or learned spatial maps that are standardly used to explain the formation of place cell representations. A randomly tuned network with non-plastic connections and feedback inhibition can account for diverse phenomena observed in experimental recordings of hippocampal cells during spatial exploration, including positional tuning, multiple place fields, mixed selectivity, and remapping. The mechanism used here is more parsimonious than mechanisms requiring specialized input connections that form through evolution, development, or learning. Of note, there are known gradients of input connections targeted to regions along the proximal-distal axis of the hippocampus that coincide with differences in CA1 place selectivity (Henriksen et al., 2010) and the intrinsic properties of CA3 cells (Sun et al., 2017), but the present work suggests these gradients are not required to generate the phenomena described here. To generate place selectivity, and the other features we have characterized (Table 1), it is sufficient that network activity is sparse with a winner-take-all dynamic, where only neurons with the strongest inputs are active in any current location. Furthermore, a role for synaptic plasticity in downstream synapses was identified for associating network representations with events, and a role for plasticity in recurrent connections was found to induce representational drift (Fig. S4) as well as stabilize place fields at specific locations (Fig. 4F) and segregate subpopulations according to specialized experience-dependent inputs (Figs. 4H).

While our results constitute a sufficiency argument, the results cannot comment on necessity. The hippocampus may indeed produce its selectivity through more sophisticated means than our findings suggest. Nonetheless, the simple mechanism proposed here accounts for a substantial number of perplexing phenomena from the empirical hippocampal literature. The ability to remap is facilitated by the broad selectivity, allowing neurons to respond uniquely to different input patterns from different environments, the resulting activity is both place specific and environment specific, as well as frame-specific (Figs. 1,2, 3E,F). Our model thus addresses the tension in existing descriptions requiring preconfigured or overly flexible connections. The success of this simplified model is in revealing how complex spatial representations can emerge from relatively simple network architectures while facilitating memory representations that are robust, flexible, and spontaneously generated in the absence of a global optimization principle or objective function.

Nonetheless, future research should explore the integration of more biologically detailed mechanisms within this simplified framework, potentially incorporating additional forms of synaptic plasticity and neuromodulation. Moreover, the role of short time-scale synchrony caused by global oscillations like local field potential (LFP) theta oscillations could be studied in terms of its effect of coordinating spatial and temporal features of the network activity. Investigating the role of different inhibitory interneuron types and their contributions to spatial coding and memory formation could provide further insights into the intricacies of hippocampal function. There is some evidence that inhibitory neurons form subpopulation-specific cell assemblies of coactivity with excitatory neurons, and these clustering effects may be relevant to versions of the DivSparse model incorporating biases to some neurons instead of others (van Dijk and Fenton, 2018; Huszar et al., 2022). Additionally, this model offers specific hypotheses for experimental validation, based on the model’s generation of environment-specific anti-cofiring cells (Levy et al., 2023), which predicts that anti-cofiring cells emerge due to long-term potentiation, and that anti-cofiring is a signature of self-organizing assemblies of co-potentiated cells.

### Network Sparsity and Computations

Multiple mechanisms may accomplish the network sparsification that we find is crucial for explaining diverse hippocampal phenomena. These mechanisms include divisive normalization, E-I coupling, input pruning and differences in intrinsic excitability. Spatial information is abundant in the non-place cell inputs arriving at hippocampus from the entorhinal and sensory cortices, allowing the hippocampus to apply a simple computation, such as the DivSparse mechanism presented in this paper (Olypher et al., 2002; Fenton, 2024). These findings provide new insights into the emergence and transformation of spatial representations in the hippocampus. By demonstrating that a randomly tuned network with simple inhibitory dynamics can account for a wide range of hippocampal phenomena, not only firing field statistics (Mainali et al., 2025), this work not only advances theoretical models of hippocampal function but also accounts for idiosyncratic and unexplained features of place cell activity. As such, this framework readily generates experimental hypotheses to probe into the neural mechanisms underlying spatial cognition and the hippocampus. For example, the model suggests that network sparsity is the key variable upon which hippocampal and place cell phenomena depends. Crucially, the results demonstrate that effective sparsity can be achieved by multiple neurobiological mechanisms such as changes in inhibition, E-I coordination, connectivity, excitability or their combination. Importantly, sparsity is not a feature of a single cell and can’t be assessed by single cell measurements, rather sparsity is a network property that depends on collective neuronal interactions.

Although the model is agnostic to the modality of input features (e.g. spatial or nonspatial), the findings support the cognitive map hypothesis, providing a mechanism for many unexplained phenomena that challenged the notion of a unitary and allocentric map (O’Keefe and Nadel, 1978). Furthermore, the model is also applicable to nonspatial task correlates of hippocampal activity like jumping, or being dropped, or informative sounds (Moita et al., 2003, 2004; Lenck-Santini et al., 2008; Aronov et al., 2017). Random input connections, acting as universal encoders, combined with sparse outputs allow the downstream network neurons to encode or recollect whatever was being presented by the inputs.

By incorporating local activity-dependent plasticity at recurrent synapses, this model also addresses the phenomena of overdispersion, representational drift, and memory tagging (Fenton and Muller, 1998; Olypher et al., 2002; Garner et al., 2012; Ramirez et al., 2013; Rule et al., 2019; Zaki et al., 2025). These features are crucial for understanding the dynamic nature of hippocampal representations and their ability to support complex behaviors and memory processes over extended periods of time. The findings on remapping and memory-associated tagging demonstrate that changes in single-cell tuning across the network can occur due to alterations in input patterns during distinct contexts rather than due to modifications of the input connections. This perspective highlights how remapping can facilitate memory associations and context-specific codes despite the arbitrary nature of the changes that experimentalists define as distinct “contexts.” In addition to reproducing the remapping phenomena, the ability of this model to reproduce the diverse findings of multiple enlarged place fields in larger environments, overdispersion, representational drift, anti-cofiring, and spatial frame ensemble preference emphasize the expressive power of broad selectivity and winner-take-all dynamics in spatial information processing.

### Conclusions and Outlook

The experimental features of the hippocampus reproduced in this work are typically associated with specialized circuits or plasticity at hippocampal synapses, which is why recent results from investigating randomly-connected networks without synaptic plasticity (Chandra et al., 2025; Khona et al., 2025; Mainali et al., 2025), may seem at odds with standard concepts of place cell firing, learning, memory, and synaptic plasticity phenomena being facets of the same underlying computational process. Nonetheless, implementations of randomly connected networks, optimized EI networks, divisive normalization and assembly-tagging have shown promising characteristics in diverse domains of machine learning and neuroscience (Dasgupta et al., 2017; Rawat et al., 2025). An important implication of the present work is that seemingly independent experimental phenomena and computational algorithms may be synthesized together in the hippocampus through its network architecture and mechanisms for firing rate homeostasis (Buzsaki et al., 2002). Aspects of the findings suggest that important properties of hippocampal spatial representations are neither pre-existing (attractor networks) or learned (associative memory) in the hippocampus but may instead be an instance of diverse inputs projected onto internally-organized and self-regulated complex dynamical systems. Accordingly, mechanisms that transform spatial representations due to behavioral relevance may therefore be studied separately from the representations themselves, as only the former can be properly considered a “learning function” that engages the hippocampus. The range of behaviors that depend on HPC function makes it difficult to leverage the optimization of a known learning objective to compare artificial and biological representations. While this may be a conceptual limitation, the present findings demonstrate that several features of hippocampal representations do not necessarily require learning or specialized connections. Our findings suggest that sparsity may be a computational objective of hippocampal function, and thus this work may provide a framework to determine other computational objectives of transforming spatial representations, as opposed to forming them.

The DivSparse model could therefore be used as the initialization of a network that is later trained on one or more hypothetical computational objectives, which may or may not be a globally-defined, information-maximizing function. In future work, we explore the possibility that the DivSparse model allows for multisensory, multi-frame representations that can be organized using local-learning principles, that minimizes metabolic demands across neuronal populations (Fenton et al., 2023), settling into input-specific assemblies that can be memory-tagged. We hypothesize that such a local-learning rule will act to regularize co-existing spatial representations that may otherwise cause interference across conflicting spatial frames as an exemplar of conflicting memories in other domains of knowledge. This self-supervised approach would rely on the pre-existing representations conditioned on the inputs, which greatly simplify the multi-frame learning and memory problem, as opposed to learning these on top of learning the spatial representations themselves. The significance of such a model for hippocampal experimentalists cannot be overstated. Biological considerations of metabolic homeostasis, energetic demands, and neuronal turnover are often abstracted away when modeling the formation of learned representations in neural network models. In contrast to this approach, a homeostatic DivSparse model would leverage biological considerations to model efficient representations that can be compared to experimental findings, bridging the neurophysiological literature on hippocampal function at the cellular level with the computational properties that emerge at the network level. We are encouraged by recent work on recurrent neural networks that can simultaneously achieve divisive normalization while solving a learning objective (Rawat et al., 2025). A self-supervised extension to this model will also be a good candidate for bridging the gap between metabolic and computational properties of a network.

In summary, the present work consolidates findings from experimental, theoretical, and modeling work into a synthesized framework that explains diverse phenomena of hippocampal spatial representations. The framework is compatible with biologically plausible spiking models that are constrained by realistic dynamical properties, as well as the highly simplified, 2-step DivSparse model in one or two dimensions. Furthermore, although no global learning objective was used in this work, it segregates hippocampal phenomena that require local LTP-like plasticity from those phenomena that can spontaneously emerge directly from the structural properties of random connections, divisive normalization, and sparsification. It is not yet understood how multisensory, multimodal tasks are learned in several parts of the brain, hippocampus included. We suggest it is likely that analogous features exist in other brain regions with their own idiosyncratic input sources leading to idiosyncratic network representations. Moreover, this current work highlights that studying task-correlated representations in any given brain region can obscure the biological implementations leading to these representations, as there are multiple solutions to the problem of generating a given representation. Specifically, stable and persistent context-specific location-specific representations were generated without changing the network. These representations are latent; they are present in the connectivity prior to the experience during which they are observed, consistent with observed preplay phenomena (Dragoi and Tonegawa, 2011, 2013). Such location-specific representations are widely-considered to be critical evidence of experience-dependent memory, and when the active cells have been identified and manipulated to elicit conditioned behavior, is taken as evidence that the cells are “engram cells,” representing the enduring change that experience has caused in a neural circuit. The present work indicates the possibility of an alternative interpretation, in which the so-call engram was preexisting and therefore neither an engram nor the result of any other change intrinsic to the neural system. This existence proof questions the validity of current interpretations of engram and neural coding phenomena, especially concerning hippocampal place cells (Josselyn and Tonegawa, 2020; Dragoi, 2024; Fenton, 2024; Mainali et al., 2025).

We have presented a model to evaluate concepts, and do not attempt to mimic the hippocampus or any other neural system. Although the model incorporates known features of the hippocampal architecture, we have taken a deliberately parsimonious approach, providing a minimalist model in both LIF and analytical forms. The model is sufficient to reproduce diverse and well-documented experimental phenomena, laying the groundwork for studying learning and memory from a sufficiently simple foundation that facilitates biologically-relevant investigations and interpretations of the observations that are yet to come from ingenious experimentalists.

## Acknowledgements

Supported by NIH grant R01MH132204 to AAF

## METHODS

### Input architecture

(See table of parameter choices below - Table 2–Table 4)

Trajectories around a one-dimensional circular environment or a two-dimensional rectangular environment were simulated to sample input activity for the network. Input neurons generate spikes using mixture of Gaussian spatial filters with Gaussian centers chosen at random locations (∼6 locations per input). For each location “pos” sampled during the simulation, a probability of firing was calculated as:

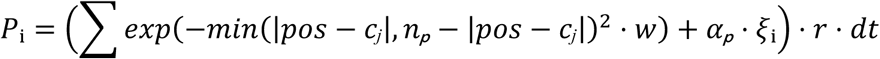

Where pos is the position at time t, cj refers to the *j*-th center of input neuron i, *n_p_* is the number of positions, while *w* specifies the width of the Gaussian envelope. The minimization step ensures that periodic circular distances are properly calculated. Noise is added to the inputs in the form of pink noise ξ_i_ modulated by scalar value *α_p_* using a pre-existing MATLAB package (https://github.com/cortex-lab/MATLAB-tools/blob/master/pinknoise.m). These input activities are passed into the LIF network with feedforward all-to-all connections and weights drawn from a scaled uniform distribution.

### Leaky integrate and fire network with inhibition

A previously published leaky integrate and fire network (Fenton et al., 2023; Levy et al., 2023) of 500 E and 50 I neurons was modified to have fixed, all-to-all EI and IE connections which were preset for the simulation. EE and II connections were set to zero, since these are not needed for place tuning in this network, while EI and IE connections were set to the same value (Fig. 1, Fig. S2) or drawn from a scaled uniform distribution (Fig S1). The activity of an E neuron in the network is modulated by the membrane potential, which changes due to three factors: 1) the sum of the neuron’s weighted presynaptic inputs, 2) the strength of the feedback inhibition, and 3) the leaky membrane potential that gradually returns to rest. The differential equation governing network dynamics is expressed as:

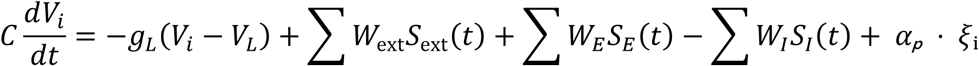

Where *V_i_* is the membrane potential of the *i-*th neuron at time t, *S*_ext_ is the spiking activity of the external inputs, while *S*_’_ and *S*_(_ refer to the spiking activities of the E and I recurrent neurons with *W* corresponding to the weights of the external input and recurrent connections. *g_L_*, *V_L_*, and *C* are biological parameters of the LIF network, determining the time constant and resting potential (See Table T2). When the membrane potential crosses the threshold (see table T2) a spike is recorded, and the potential is returned to the resting potential. The parameters of the network were simplified to C=gL=threshold=1 for figures 1 and 3, with the resting potential set to zero, this parametrization is called the Slow LIF model. For the Fast LIF model in Figure 3, realistic biological parameters were chosen as shown in table T2.

To compensate for the inhibiting effects of the fast membrane time constant in the biologically plausible model of Figure 1H-J, the rate and width of the input spiking generation (*r* and *w*^-1^) were increased while the input, EI, and IE connections were scaled down by Scale_Input_, Scale_EI_, and Scale_IE_, respectively to avoid the saturation of activities:

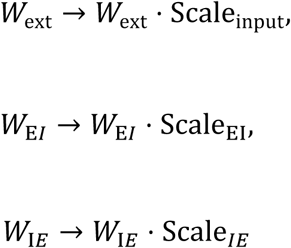

All simulations were performed using the Euler method with distinct integration time steps for the Fast and Slow LIF models (Table T.2)

### Assembly Tagging decoder and remapping analysis

Since memory encoding in the HPC appears to be subpopulation-specific (like memory-tagged and reactivated “engram” cells), a decoder was used to decode locations from the activations of E neuron subpopulations. The first half of the simulation (Train) is used to determine the location of highest trial-averaged activity for each neuron, and these locations are used in the next half (Test) as “votes” for the decoder. Neurons that are inactive do not vote, and the rest contribute one vote per neuron, registered as the highest-activity position in the first half. For a given interval of time (1s), if a neuron is active, it casts its vote into the decoder. The location with the largest number of votes is selected as the decoded position for that time point.

Mathematically, the assembly tagging decoder can be summarized in the following manner. To compute *s_p_(t)*, the decoded stimulus at a given position *p*, at time *t* is:

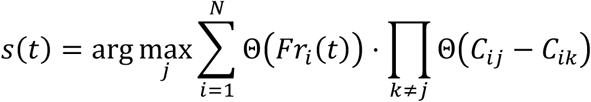

Where Fr_i_(t) is the activity of neuron i at time t. C_ij_ is a Neuron-by-Position matrix specifying the session-averaged activity of neuron i at position j during a training simulation. Θ is the Heaviside step function equal to 1 if its argument is positive and 0 otherwise. Thus, the first Heavyside function ensures only active neurons are counted, while the second Heavyside function ensures a vote for position “k” is only counted if that is the position of highest activity for neuron i. If not, the entire term is canceled by negative values in ∏*_k≠j_* Θ(*C_ij_* − *C_ik_*). Furthermore, the position “j” with the most votes across N neurons is the decoded position s(t). The assembly tagging decoder encourages a sparse output as well as selective subpopulations that are dominant in their respective location.

### Analysis of effective tuning from random inputs

The effect of the random spatial filters and input connections on the output E neurons was computed before the effects membrane potential dynamics and feedback inhibition to understand the effective role of the LIF network (Fig. 1B). A matrix **T** representing the filters of the input neurons and positions was generated with random binary entries where the 1s refer to centers of the spatial filters for input neurons. Each neuron had ∼6 centers in a multi-peaked mixture of input selectivity (20% of locations). Multiplying this matrix by the random input connection matrix **U** yields the feedforward linear activations for each output neuron. Normalizing each column across the neurons generates an effective tuning matrix (ETM).

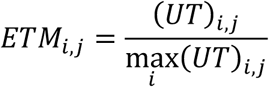

For each position, the ETM encodes a value near 1 for the subpopulation that is most likely to dominate the winner-take-all dynamics at that location. Position-averaged membrane potentials were also computed after running the LIF simulation, sorted along neurons by location preference, and used in Fig 1. to compare with the distribution of the network’s effective tuning.

### Contextual and positional associations in LIF model

Simulations for the tagging mechanism were split between training and testing simulations. During training simulations, a simulation was performed to extract contextual and positional associations. Positional associations were generated by including a special location in the trajectories that model a shock zone, introducing a special shock signal that will generate associations as follows. Whenever the simulated trajectory crosses the special location, the shock signal causes the input activity to be increased, but only for inputs with filters that overlap the special location. At the end of the training simulation, the neurons that cross an activity threshold are tagged by a downstream readout that associates this sparse subset of activations with an avoidance response (i.e. “turn around”). The ability of the LIF model to generate selective outputs causes the special increase in input activity to be expressed in subpopulations that are easy to tag and associate. Briefly, after tagging, whenever the activity of 70% of tagged cells exceeds a baseline, the downstream reader triggers a reversal of the current direction in the simulated trajectories. Therefore, reactivation of a tagged subpopulation leads to avoidance behavior in the model.

To program position-independent associations of context, the primacy of neurons with respect to their first 20 spikes was computed. A primacy set refers to the set of neurons that most rapidly reach their first n-spikes (Wilson et al., 2017). This is useful since different input patterns from distinct contexts generate different primacy sets, and many cells fail to reach 20 spikes for a given context. The assemblies used for contextual associations were grouped together according to primacy, grouping together neurons with similar primacy (Fig. S5 A, B). Each primacy set was associated to a particular downstream neuron, which is only active only when 90% of the tagged assembly is active. When this occurs, trajectories freeze for a moment before resuming in the same direction. During the testing stage, only the associations between the output E neurons and the downstream readout’s activity is used to recall a general context via freezing or a special location via avoidance using the same population of cells and different contexts for each association (Fig S6).

### Spike-timing-dependent plasticity (STDP) in the EI model

STDP rules alter EI and IE connections depending on the relative timing of spikes between presynaptic and postsynaptic neurons in each connection as done in (Levy et al 2023). The eligibility traces of excitatory or inhibitory neurons *x* decays exponentially and increases when a spike occurs. This can be expressed in the following form:

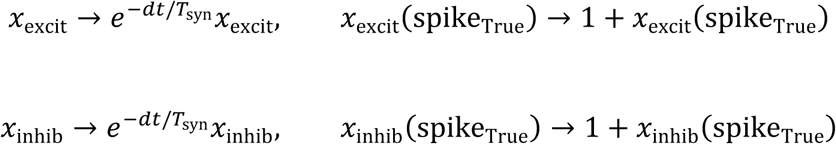

From these eligibility traces, asymmetric STDP and symmetric STDP is applied to EI and IE synapses, respectively, by combining the with a binary vector of active and inactive neurons:

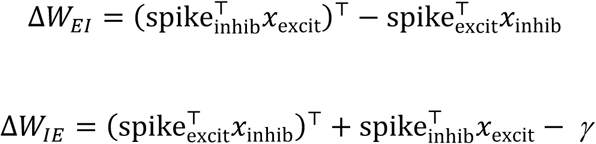

Where “spike_excit_” (or “spike_inhib_”) denotes the binary vector of excitatory (or inhibitory) neurons that were active at a given time. *ψ* is a constant that defines the zero-point weight change, thus setting the temporal delay necessary to increase or decrease the IE synapse strength. The resulting weight changes can subsequently be scaled by a learning rate (η).

### Simplified Positional Network, 1D Case

The Effective Tuning Matrix (ETM) used to analyze the feedforward effect of the random filters and random connections was also used in the simplifying model to generate activities. In the simple model, the ETM is generated as explained above. To model the effect of the EI network, which includes the feedback inhibition, threshold spiking, and leaky membrane potentials, the ETM is sparsified by a threshold that set all smaller values to zero.

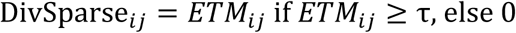

The resulting tuning matrix, DivSparse, is now sparse and selective. Each column contains a number near 1 for the cell-location pairs that represent the center of a place field. Next, temporal dynamics are drawn from this matrix using the same method of spike generations as for the input neurons, using discrete inhomogeneous Poisson spikes sampling from the DivSparse matrix directly, with added homogeneous Poisson background noise to generate the results of figure 2.

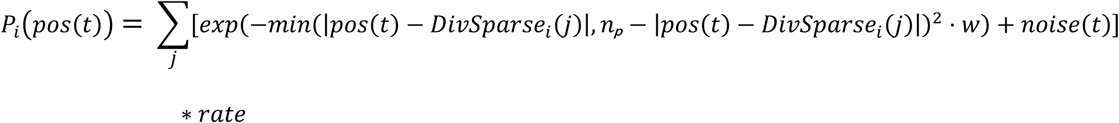

Therefore, DivSparse represents the network output of E neurons directly, after abstracting EI model effects, but prior to sampling specific inhomogeneous Poisson spikes in time.

The DivSparse model is naturally converted into a rate model of activations by sampling rates directly from the DivSparse matrix at a given location. The rate-based version of the model was used in figures 3 and 4. In figure 3, the different DivSparse models were designed using different assumptions on the structure of the input neuron tuning that defines the Effective Tuning Matrix (ETM) prior to thresholding. The first version ignored the divisive normalization step. The second was the same model described above, with uniform random input connections, and tuning generated from a mixture of delta functions (0s and 1s). The last version uses a mixture of gaussian tuning, replacing the 1s in the mixture of delta tuning with the center of a gaussian bump, therefore adding signal correlations to the activities occurring in nearby locations.

### Positional Information (Ipos)

Positional information (Ipos) for a given cell was calculated according to (Kelemen and Fenton 2010). Briefly, probabilities of having activity n spikes constrained by position are compared to unconstrained spiking probabilities as shown in this equaiton:

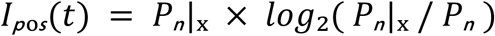

At a given time t, the probability of firing at x(t) is compared to the unconstrained probability to determine the positional information for a given cell at time t. The ensemble Ipos (Fig 3) was computed by summing frame-specific Ipos scores across the population.

### Simplified Positional Network, 2D Case, Plasticity, and LTP

For 2D simulations, the ETM’s were computed in the same manner, except the matrix is now a 3D tensor representing neurons, x-dimension, and y-dimension. This ETM is also computed from random filters and connections, except now the random filters are 2D binary matrices for each neuron instead of 1D vectors. Thus, each slice of the 3D ETM tensor along the neuron axis is a particular neuron’s feedforward spatial map, with the entire neuronal population normalized along each location (pixel) and thresholded in the same manner as the 1D case. For each neuronal slice of the resulting 3D tensor, 2D Gaussian envelopes are drawn centered at the remaining nonzero value after sparsification. The peak height of each Gaussian envelope is determined by the nonzero value, and the width *w* is drawn for each input from a lognormal distribution (See Table T4). The resulting Gaussian mixture using the DivSparse matrix corresponds to the rate model of positional activities that will be sampled by the simulated trajectories. To incorporate a simple form of STDP into the model, recurrent weights were initialized at zero and set to decay to zero with a given preset time constant (Table T4). For a given sliding time window of length T_Learning_, neurons *i* active above a *Thresh_Spike_* value change connections with neurons *k* active later. *W_i,k_* weights increase while *W_k,i_* decrease. The magnitude of these changes depend on the time *j* relative to neuron *i*’s activity and is given by the following equations:

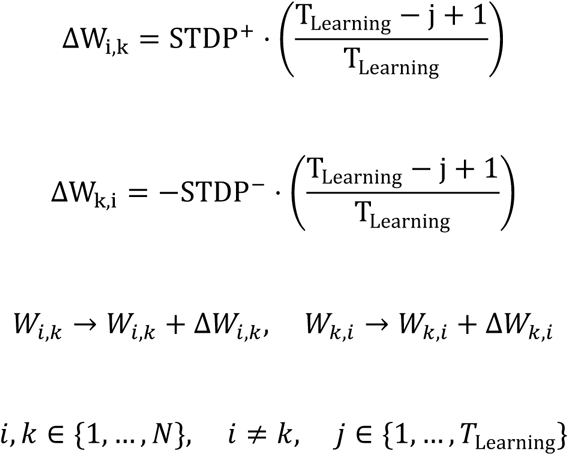

Once all the connections for neurons active at the first time point are accounted for, the T_Learning_ window slides one time step and the STDP process repeats. Therefore, neurons that are active earlier influence neurons that tend to be active later. This local plasticity rule is meant to model short term effects of STDP in a simple rate-based model to test hypotheses about transforming spatial maps. Lastly, to model long term plasticity (LTP) associations in the network, a special location was generated to represent a shock zone. When the trajectories cross the shock zone, all neurons that are above the threshold (Threshold_LTP_) increase recurrent connections with each other (*W_i,k_* = 2) and maintain these connections for the rest of the simulation. Note these are the unnormalized weights, and these are normalized moment-to-moment by max-normalizing ([0, 1]). During new contexts, input filters are shuffled, and new special locations are generated to test the effect of remapping on plasticity and LTP.

### Place Cell Metrics

<u>Overdispersion statistics</u> (Fig. S8) were calculated for all positionally selective cells at their respective place field location. The activity resulting from multiple passes through a place field center were computed after low firing activity was filtered out (less than 80% of the peak), while a random choice was made for cells with multiple place fields. The resulting distribution was Z-Scored, showing the dispersion of firing rate activities at the center of the respective field, allowing for comparisons between plastic and aplastic connections in the model. <u>Map stability</u> is defined as the Pearson correlation of positional tuning at time t compared to the positional tuning at the final time step. <u>Anti-cofiring statistics</u> are determined using a Pearson correlation of the neural activities after down sampling by averaging every 10 neighboring time points. A pairwise correlation is considered anti-cofiring if the pairwise correlation value is below -0.05, and a particular cell’s anti-cofiring power is equal to the proportion of anti-cofiring pairs across all synaptic partners.

**Table T2:**
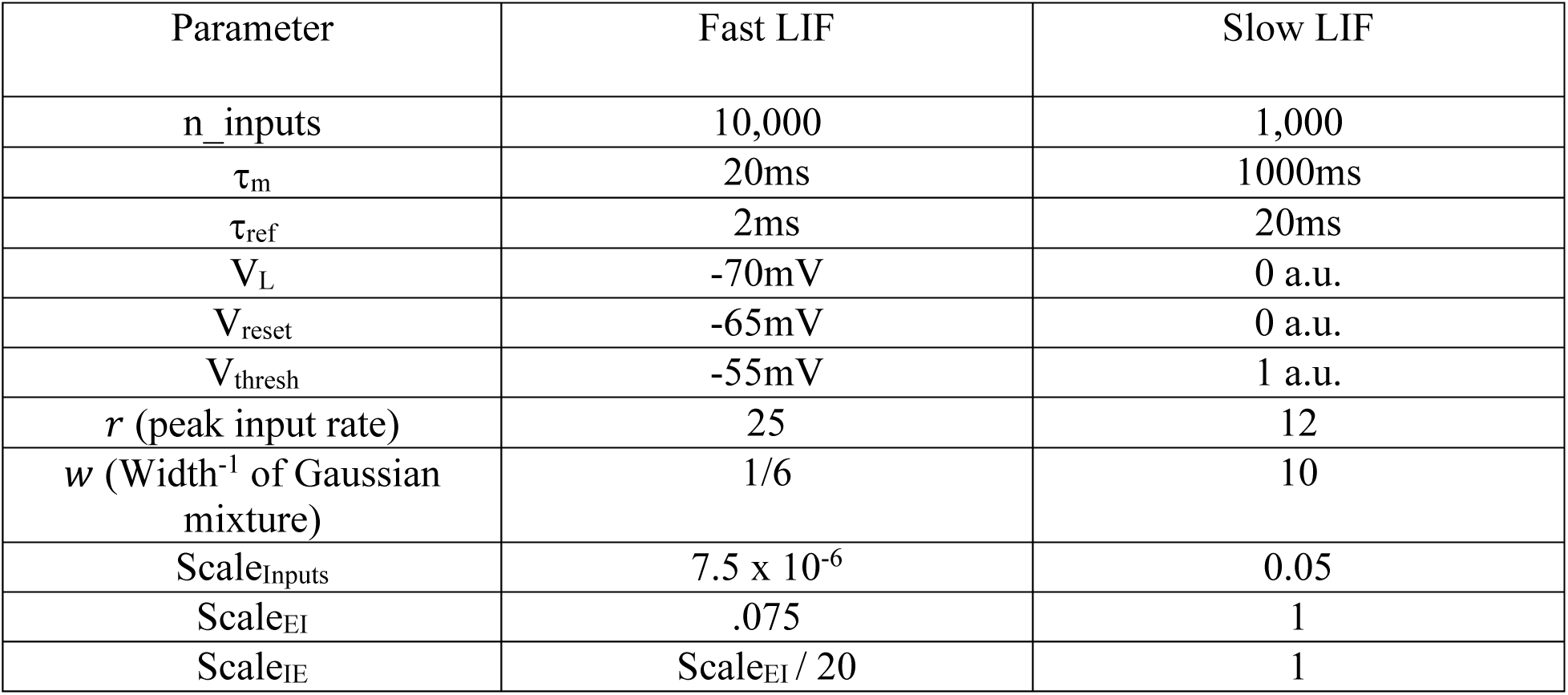
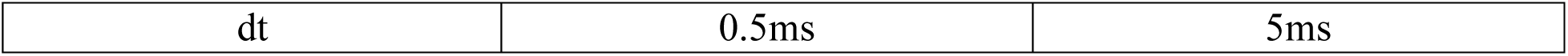
Fixed Weight EI Model Parameters.

| Parameter | Fast LIF | Slow LIF |
| --- | --- | --- |
| $n_{\text{inputs}}$ | 10,000 | 1,000 |
| $\tau_m$ | 20ms | 1000ms |
| $\tau_{\text{ref}}$ | 2ms | 20ms |
| $V_L$ | -70mV | 0 a.u. |
| $V_{\text{reset}}$ | -65mV | 0 a.u. |
| $V_{\text{thresh}}$ | -55mV | 1 a.u. |
| $r$ (peak input rate) | 25 | 12 |
| $w$ (Width<sup>-1</sup> of Gaussian mixture) | 1/6 | 10 |
| $\text{Scale}_{\text{Inputs}}$ | $7.5 \times 10^{-6}$ | 0.05 |
| $\text{Scale}_{\text{EI}}$ | .075 | 1 |
| $\text{Scale}_{\text{IE}}$ | $\text{Scale}_{\text{EI}} / 20$ | 1 |
| dt | 0.5ms | 5ms |

**Table T3:** STDP Model Parameters.

| Parameter | Value |
| --- | --- |
| $\tau_{\text{syn}}$ | 20ms |
| $\gamma$ | 0.0001 |
| $W_{\text{Max}}$ | 0.5 |
| $W_{\text{Initial\_Low}}$ | 0.005 |
| $W_{\text{Initial\_High}}$ | 0.25 |
| $\eta$ | 0.0005 |

**Table T4:** 2D Simplified Model Parameters.

| Parameter | Values |
| --- | --- |
| n_inputs | 300 |
| n_excit | 100 |
| $r$ (mean peak rate) | 10 |
| $w$ (Scale<sup>-1</sup> of lognormal mixture of rates) | 75 |
| Thresh<sub>Spike</sub> | 0.35 |
| T<sub>Learning</sub> | 15 Time-bin windows |
| $W_{\text{EE\_Initial}}$ | 0 |
| $W_{\text{EE\_Decay}}$ | 35% Per Time Bin |
| Sparsity<sub>EE</sub> | 50% Connections |
| Threshold<sub>LTP</sub> | 0.9 |
| STDP+ | 0.1 |
| STDP- | 1 |
| Max Weight (LTP before norm) | 2 |

## REFERENCES

Aronov D, Nevers R, Tank DW (2017) Mapping of a non-spatial dimension by the hippocampal-entorhinal circuit. Nature 543:719–722.

Barry C, Burgess N (2007) Learning in a geometric model of place cell firing. Hippocampus 17:786–800.

Barry C, Lever C, Hayman R, Hartley T, Burton S, O’Keefe J, Jeffery K, Burgess N (2006) The boundary vector cell model of place cell firing and spatial memory. Rev Neurosci 17:71–97.

Barry JM, Rivard B, Fox SE, Fenton AA, Sacktor TC, Muller RU (2012) Inhibition of protein kinase Mzeta disrupts the stable spatial discharge of hippocampal place cells in a familiar environment. J Neurosci 32:13753–13762.

Beggs JM, Plenz D (2003) Neuronal avalanches in neocortical circuits. J Neurosci 23:11167–11177.

Bernard C, Wheal HV (1994) Model of local connectivity patterns in CA3 and CA1 areas of the hippocampus. Hippocampus 4:497–529.

Bittner KC, Milstein AD, Grienberger C, Romani S, Magee JC (2017) Behavioral time scale synaptic plasticity underlies CA1 place fields. Science 357:1033–1036.

Bittner KC, Grienberger C, Vaidya SP, Milstein AD, Macklin JJ, Suh J, Tonegawa S, Magee JC (2015) Conjunctive input processing drives feature selectivity in hippocampal CA1 neurons. Nat Neurosci 18:1133–1142.

Bliss TV, Gardner-Medwin AR (1973) Long-lasting potentiation of synaptic transmission in the dentate area of the unanaestetized rabbit following stimulation of the perforant path. J Physiol 232:357–374.

Bliss TV, Collingridge GL (1993) A synaptic model of memory: long-term potentiation in the hippocampus. Nature 361:31–39.

Blum KI, Abbott LF (1996) A model of spatial map formation in the hippocampus of the rat. Neural Comput 8:85–93.

Brown EN, Frank LM, Tang D, Quirk MC, Wilson MA (1998) A statistical paradigm for neural spike train decoding applied to position prediction from ensemble firing patterns of rat hippocampal place cells. J Neurosci 18:7411–7425.

Brun VH, Ytterbo K, Morris RG, Moser MB, Moser EI (2001) Retrograde amnesia for spatial memory induced by NMDA receptor-mediated long-term potentiation. J Neurosci 21:356–362.

Brun VH, Otnass MK, Molden S, Steffenach HA, Witter MP, Moser MB, Moser EI (2002) Place cells and place recognition maintained by direct entorhinal-hippocampal circuitry. Science 296:2243–2246.

Buzsaki G (1984) Feed-forward inhibition in the hippocampal formation. Prog Neurobiol 22:131–153.

Buzsaki G, Kaila K, Raichle M (2007) Inhibition and brain work. Neuron 56:771–783.

Buzsaki G, Csicsvari J, Dragoi G, Harris K, Henze D, Hirase H (2002) Homeostatic maintenance of neuronal excitability by burst discharges in vivo. Cereb Cortex 12:893–899.

Chandra S, Sharma S, Chaudhuri R, Fiete I (2025) Episodic and associative memory from spatial scaffolds in the hippocampus. Nature 638:739–751.

Chettih SN, Mackevicius EL, Hale S, Aronov D (2024) Barcoding of episodic memories in the hippocampus of a food-caching bird. Cell 187:1922–1935 e1920.

Cho YH, Giese KP, Tanila H, Silva AJ, Eichenbaum H (1998) Abnormal hippocampal spatial representations in alphaCaMKIIT286A and CREBalphaDelta-mice. Science 279:867–869.

Chung A, Jou C, Grau-Perales A, Levy ERJ, Dvorak D, Hussain N, Fenton AA (2021) Cognitive control persistently enhances hippocampal information processing. Nature 600:484–488.

DasGupta S, Waddell S (2008) Learned odor discrimination in Drosophila without combinatorial odor maps in the antennal lobe. Curr Biol 18:1668–1674.

Dasgupta S, Stevens CF, Navlakha S (2017) A neural algorithm for a fundamental computing problem. Science 358:793–796.

Dayan P (1993) Improving Generalization for Temporal Difference Learning: The Successor Representation. Neural Computation 5:613–624.

de Cothi W, Barry C (2020) Neurobiological successor features for spatial navigation. Hippocampus 30:1347–1355.

de Snoo ML, Miller AMP, Ramsaran AI, Josselyn SA, Frankland PW (2023) Exercise accelerates place cell representational drift. Current Biology 33:R96–R97.

Dobrunz LE, Stevens CF (1999) Response of hippocampal synapses to natural stimulation patterns. Neuron 22:157–166.

Dragoi G (2024) The generative grammar of the brain: a critique of internally generated representations. Nat Rev Neurosci 25:60–75.

Dragoi G, Tonegawa S (2011) Preplay of future place cell sequences by hippocampal cellular assemblies. Nature 469:397–401.

Dragoi G, Tonegawa S (2013) Distinct preplay of multiple novel spatial experiences in the rat. Proc Natl Acad Sci U S A 110:9100–9105.

Dupret D, O’Neill J, Pleydell-Bouverie B, Csicsvari J (2010) The reorganization and reactivation of hippocampal maps predict spatial memory performance. Nat Neurosci 13:995–1002.

Epsztein J, Brecht M, Lee AK (2011) Intracellular determinants of hippocampal CA1 place and silent cell activity in a novel environment. Neuron 70:109–120.

Fenton AA (2024) Remapping revisited: how the hippocampus represents different spaces. Nature Reviews Neuroscience.

Fenton AA, Muller RU (1998) Place cell discharge is extremely variable during individual passes of the rat through the firing field. Proc Natl Acad Sci U S A 95:3182–3187.

Fenton AA, Csizmadia G, Muller RU (2000a) Conjoint control of hippocampal place cell firing by two visual stimuli. II. A vector-field theory that predicts modifications of the representation of the environment. J Gen Physiol 116:211–221.

Fenton AA, Csizmadia G, Muller RU (2000b) Conjoint control of hippocampal place cell firing by two visual stimuli. I. The effects of moving the stimuli on firing field positions. J Gen Physiol 116:191–209.

Fenton AA, Wesierska M, Kaminsky Y, Bures J (1998) Both here and there: simultaneous expression of autonomous spatial memories in rats. Proc Natl Acad Sci U S A 95:11493–11498.

Fenton AA, Hurtado JR, Broek JAC, Park E, Mishra B (2023) Do Place Cells Dream of Deceptive Moves in a Signaling Game? Neuroscience 529:129–147.

Fenton AA, Kao H-Y, Neymotin SA, Olypher AV, Vayntrub Y, Lytton WW, Ludvig N (2008) Unmasking the CA1 ensemble place code by exposures to small and large environments: more place cells and multiple, irregularly-arranged, and expanded place fields in the larger space. J Neurosci 28:11250–11262.

Fenton AA, Lytton WW, Barry JM, Lenck-Santini PP, Zinyuk LE, Kubik S, Bures J, Poucet B, Muller RU, Olypher AV (2010) Attention-like modulation of hippocampus place cell discharge. J Neurosci 30:4613–4625.

Garner AR, Rowland DC, Hwang SY, Baumgaertel K, Roth BL, Kentros C, Mayford M (2012) Generation of a synthetic memory trace. Science 335:1513–1516.

Gershman SJ, Moore CD, Todd MT, Norman KA, Sederberg PB (2012) The successor representation and temporal context. Neural Comput 24:1553–1568.

Geva N, Deitch D, Rubin A, Ziv Y (2023) Time and experience differentially affect distinct aspects of hippocampal representational drift. Neuron 111:2357–2366.e2355.

Gill JV, Lerman GM, Zhao H, Stetler BJ, Rinberg D, Shoham S (2020) Precise Holographic Manipulation of Olfactory Circuits Reveals Coding Features Determining Perceptual Detection. Neuron 108:382–393.e385.

Gothard KM, Skaggs WE, McNaughton BL (1996a) Dynamics of mismatch correction in the hippocampal ensemble code for space: interaction between path integration and environmental cues. J Neurosci 16:8027–8040.

Gothard KM, Skaggs WE, Moore KM, McNaughton BL (1996b) Binding of hippocampal CA1 neural activity to multiple reference frames in a landmark-based navigation task. J Neurosci 16:823–835.

Grienberger C, Milstein AD, Bittner KC, Romani S, Magee JC (2017) Inhibitory suppression of heterogeneously tuned excitation enhances spatial coding in CA1 place cells. Nat Neurosci 20:417–426.

Harland B, Contreras M, Souder M, Fellous JM (2021) Dorsal CA1 hippocampal place cells form a multi-scale representation of megaspace. Curr Biol.

Hartley T, Burgess N, Lever C, Cacucci F, O’Keefe J (2000) Modeling place fields in terms of the cortical inputs to the hippocampus. Hippocampus 10:369–379.

Hayashi Y (2019) NMDA Receptor-Dependent Dynamics of Hippocampal Place Cell Ensembles. The Journal of Neuroscience 39:5173.

Henriksen EJ, Colgin LL, Barnes CA, Witter MP, Moser MB, Moser EI (2010) Spatial representation along the proximodistal axis of CA1. Neuron 68:127–137.

Hok V, Chah E, Save E, Poucet B (2013) Prefrontal cortex focally modulates hippocampal place cell firing patterns. J Neurosci 33:3443–3451.

Hok V, Lenck-Santini PP, Roux S, Save E, Muller RU, Poucet B (2007) Goal-related activity in hippocampal place cells. J Neurosci 27:472–482.

Hollup SA, Molden S, Donnett JG, Moser MB, Moser EI (2001) Accumulation of hippocampal place fields at the goal location in an annular watermaze task. J Neurosci 21:1635–1644.

Huszar R, Zhang Y, Blockus H, Buzsaki G (2022) Preconfigured dynamics in the hippocampus are guided by embryonic birthdate and rate of neurogenesis. Nat Neurosci 25:1201–1212.

Isaac JT, Buchanan KA, Muller RU, Mellor JR (2009) Hippocampal place cell firing patterns can induce long-term synaptic plasticity in vitro. J Neurosci 29:6840–6850.

Jackson J, Redish AD (2007) Network dynamics of hippocampal cell-assemblies resemble multiple spatial maps within single tasks. Hippocampus 17:1209–1229.

Josselyn SA, Tonegawa S (2020) Memory engrams: Recalling the past and imagining the future. Science 367.

Kanter BR, Lykken CM, Avesar D, Weible A, Dickinson J, Dunn B, Borgesius NZ, Roudi Y, Kentros CG (2017) A Novel Mechanism for the Grid-to-Place Cell Transformation Revealed by Transgenic Depolarization of Medial Entorhinal Cortex Layer II. Neuron 93:1480–1492 e1486.

Kelemen E, Fenton AA (2010) Dynamic grouping of hippocampal neural activity during cognitive control of two spatial frames. PLoS Biol 8:e1000403.

Kelemen E, Fenton AA (2016) Coordinating different representations in the hippocampus. Neurobiol Learn Mem 129:50–59.

Kentros C (2006) Hippocampal place cells: The “where” of episodic memory? Hippocampus 16:743–754.

Kentros C, Hargreaves E, Hawkins RD, Kandel ER, Shapiro M, Muller RV (1998) Abolition of long-term stability of new hippocampal place cell maps by NMDA receptor blockade. Science 280:2121–2126.

Kentros CG, Agnihotri NT, Streater S, Hawkins RD, Kandel ER (2004) Increased attention to spatial context increases both place field stability and spatial memory. Neuron 42:283–295.

Khona M, Chandra S, Fiete I (2025) Global modules robustly emerge from local interactions and smooth gradients. Nature 640:155–164.

Kubie JL, Muller RU (1991) Multiple representations in the hippocampus. Hippocampus 1:240–242.

Kubie JL, Muller RU, Bostock E (1990) Spatial firing properties of hippocampal theta cells. J Neurosci 10:1110–1123.

Laughlin SB, de Ruyter van Steveninck RR, Anderson JC (1998) The metabolic cost of neural information. Nat Neurosci 1:36–41.

Lenck-Santini PP, Fenton AA, Muller RU (2008) Discharge properties of hippocampal neurons during performance of a jump avoidance task. J Neurosci 28:6773–6786.

Lenck-Santini PP, Muller RU, Save E, Poucet B (2002) Relationships between place cell firing fields and navigational decisions by rats. J Neurosci 22:9035–9047.

Lenck-Santini PP, Rivard B, Muller RU, Poucet B (2005) Study of CA1 place cell activity and exploratory behavior following spatial and nonspatial changes in the environment. Hippocampus 15:356–369.

Lever C, Burgess N, Cacucci F, Hartley T, O’Keefe J (2002) What can the hippocampal representation of environmental geometry tell us about Hebbian learning? Biol Cybern 87:356–372.

Levy ERJ, Carrillo-Segura S, Park EH, Redman WT, Hurtado JR, Chung S, Fenton AA (2023) A manifold neural population code for space in hippocampal coactivity dynamics independent of place fields. Cell Reports 42.

Losonczy A, Zemelman BV, Vaziri A, Magee JC (2010) Network mechanisms of theta related neuronal activity in hippocampal CA1 pyramidal neurons. Nat Neurosci 13:967–972.

Mainali N, Azeredo da Silveira R, Burak Y (2025) Universal statistics of hippocampal place fields across species and dimensionalities. Neuron 113:1110–1120 e1113.

Markus EJ, Qin YL, Leonard B, Skaggs WE, McNaughton BL, Barnes CA (1995) Interactions between location and task affect the spatial and directional firing of hippocampal neurons. J Neurosci 15:7079–7094.

Marshall L, Henze DA, Hirase H, Leinekugel X, Dragoi G, Buzsaki G (2002) Hippocampal pyramidal cell-interneuron spike transmission is frequency dependent and responsible for place modulation of interneuron discharge. J Neurosci 22:RC197.

McHugh TJ, Blum KI, Tsien JZ, Tonegawa S, Wilson MA (1996) Impaired hippocampal representation of space in CA1-specific NMDAR1 knockout mice. Cell 87:1339–1349.

McHugh TJ, Jones MW, Quinn JJ, Balthasar N, Coppari R, Elmquist JK, Lowell BB, Fanselow MS, Wilson MA, Tonegawa S (2007) Dentate gyrus NMDA receptors mediate rapid pattern separation in the hippocampal network. Science 317:94–99.

Mehta MR, Barnes CA, McNaughton BL (1997) Experience-dependent, asymmetric expansion of hippocampal place fields. Proc Natl Acad Sci U S A 94:8918–8921.

Mehta MR, Quirk MC, Wilson MA (2000) Experience-dependent asymmetric shape of hippocampal receptive fields. Neuron 25:707–715.

Mizumori SJY (2006) Hippocampal place fields: A neural code for episodic memory? Hippocampus 16:685–690.

Mizumori SJY, Smith DM, Puryear CB (2007) Chapter 5 - Mnemonic contributions of hippocampal place cells. In: Neurobiology of Learning and Memory (Second Edition) (Kesner RP, Martinez JL, eds), pp 155–189. Burlington: Academic Press.

Moita MA, Rosis S, Zhou Y, LeDoux JE, Blair HT (2003) Hippocampal place cells acquire location-specific responses to the conditioned stimulus during auditory fear conditioning. Neuron 37:485–497.

Moita MA, Rosis S, Zhou Y, LeDoux JE, Blair HT (2004) Putting fear in its place: remapping of hippocampal place cells during fear conditioning. J Neurosci 24:7015–7023.

Muller R (1996) A quarter of a century of place cells. Neuron 17:813–822.

Muller RU, Stead M, Pach J (1996) The hippocampus as a cognitive graph. J Gen Physiol 107:663–694.

Muzzio IA, Levita L, Kulkarni J, Monaco J, Kentros C, Stead M, Abbott LF, Kandel ER (2009) Attention enhances the retrieval and stability of visuospatial and olfactory representations in the dorsal hippocampus. PLoS Biol 7:e1000140.

Nakazawa K, McHugh TJ, Wilson MA, Tonegawa S (2004) NMDA receptors, place cells and hippocampal spatial memory. Nat Rev Neurosci 5:361–372.

Nakazawa K, Sun LD, Quirk MC, Rondi-Reig L, Wilson MA, Tonegawa S (2003) Hippocampal CA3 NMDA receptors are crucial for memory acquisition of one-time experience. Neuron 38:305–315.

O’Keefe J (1976) Place units in the hippocampus of the freely moving rat. Exp Neurol 51:78–109.

O’Keefe J, Dostrovsky J (1971) The hippocampus as a spatial map. Preliminary evidence from unit activity in the freely-moving rat. Brain Res 34:171–175.

O’Keefe J, Nadel L (1978) The hippocampus as a cognitive map. Oxford: Clarendon Press.

O’Reilly KC, Perica MI, Fenton AA (2018) Synaptic plasticity/dysplasticity, process memory and item memory in rodent models of mental dysfunction. Schizophr Res.

Olypher AV, Lansky P, Fenton AA (2002) Properties of the extra-positional signal in hippocampal place cell discharge derived from the overdispersion in location-specific firing. Neuroscience 111:553–566.

Park E, Dvorak D, Fenton AA (2011) Ensemble Place Codes in Hippocampus: CA1, CA3, and Dentate Gyrus Place Cells Have Multiple Place Fields in Large Environments. PLoS ONE 6:e22349.

Pehlevan C, Sompolinsky H (2014) Selectivity and Sparseness in Randomly Connected Balanced Networks. PLOS ONE 9:e89992.

Pettit NL, Yuan XC, Harvey CD (2022) Hippocampal place codes are gated by behavioral engagement. Nature Neuroscience.

Pignatelli M, Ryan TJ, Roy DS, Lovett C, Smith LM, Muralidhar S, Tonegawa S (2019) Engram Cell Excitability State Determines the Efficacy of Memory Retrieval. Neuron 101:274–284 e275.

Ramirez S, Liu X, Lin PA, Suh J, Pignatelli M, Redondo RL, Ryan TJ, Tonegawa S (2013) Creating a false memory in the hippocampus. Science 341:387–391.

Rawat S, Heeger DJ, Martiniani S (2025) Unconditional stability of a recurrent neural circuit implementing divisive normalization. ArXiv.

Rich PD, Liaw HP, Lee AK (2014) Place cells. Large environments reveal the statistical structure governing hippocampal representations. Science 345:814–817.

Robinson NTM, Priestley JB, Rueckemann JW, Garcia AD, Smeglin VA, Marino FA, Eichenbaum H (2017) Medial Entorhinal Cortex Selectively Supports Temporal Coding by Hippocampal Neurons. Neuron 94:677–688 e676.

Rotenberg A, Mayford M, Hawkins RD, Kandel ER, Muller RU (1996) Mice expressing activated CaMKII lack low frequency LTP and do not form stable place cells in the CA1 region of the hippocampus. Cell 87:1351–1361.

Rotenberg A, Abel T, Hawkins RD, Kandel ER, Muller RU (2000) Parallel instabilities of long-term potentiation, place cells, and learning caused by decreased protein kinase A activity. J Neurosci 20:8096–8102.

Royer S, Zemelman BV, Losonczy A, Kim J, Chance F, Magee JC, Buzsaki G (2012) Control of timing, rate and bursts of hippocampal place cells by dendritic and somatic inhibition. Nat Neurosci 15:769–775.

Rule ME, O’Leary T, Harvey CD (2019) Causes and consequences of representational drift. Curr Opin Neurobiol 58:141–147.

Ryan TJ, Roy DS, Pignatelli M, Arons A, Tonegawa S (2015) Memory. Engram cells retain memory under retrograde amnesia. Science 348:1007–1013.

Samsonovich A, McNaughton BL (1997) Path integration and cognitive mapping in a continuous attractor neural network model. J Neurosci 17:5900–5920.

Sanzeni A, Akitake B, Goldbach HC, Leedy CE, Brunel N, Histed MH (2020) Inhibition stabilization is a widespread property of cortical networks. Elife 9.

Sanzeni A, Palmigiano A, Nguyen TH, Luo J, Nassi JJ, Reynolds JH, Histed MH, Miller KD, Brunel N (2023) Mechanisms underlying reshuffling of visual responses by optogenetic stimulation in mice and monkeys. Neuron 111:4102–4115 e4109.

Schlesiger MI, Boublil BL, Hales JB, Leutgeb JK, Leutgeb S (2018) Hippocampal Global Remapping Can Occur without Input from the Medial Entorhinal Cortex. Cell Rep 22:3152–3159.

Semon R (1921) The Mneme. London: George Allen & Unwin.

Smith DM, Bulkin DA (2014) The form and function of hippocampal context representations. Neuroscience & Biobehavioral Reviews 40:52–61.

Solstad T, Moser EI, Einevoll GT (2006) From grid cells to place cells: a mathematical model. Hippocampus 16:1026–1031.

Stachenfeld KL, Botvinick MM, Gershman SJ (2017) The hippocampus as a predictive map. Nat Neurosci 20:1643–1653.

Stefanini F, Kushnir L, Jimenez JC, Jennings JH, Woods NI, Stuber GD, Kheirbek MA, Hen R, Fusi S (2020) A Distributed Neural Code in the Dentate Gyrus and in CA1. Neuron 107:703–716 e704.

Sun Q, Sotayo A, Cazzulino AS, Snyder AM, Denny CA, Siegelbaum SA (2017) Proximodistal Heterogeneity of Hippocampal CA3 Pyramidal Neuron Intrinsic Properties, Connectivity, and Reactivation during Memory Recall. Neuron 95:656–672 e653.

Talbot ZN, Sparks FT, Dvorak D, Curran BM, Alarcon JM, Fenton AA (2018) Normal CA1 Place Fields but Discoordinated Network Discharge in a Fmr1-Null Mouse Model of Fragile X Syndrome. Neuron 97:684–697.

Thompson LT, Best PJ (1989) Place cells and silent cells in the hippocampus of freely-behaving rats. J Neurosci 9:2382–2390.

Tonegawa S, Liu X, Ramirez S, Redondo R (2015) Memory Engram Cells Have Come of Age. Neuron 87:918–931.

Tonegawa S, Tsien JZ, McHugh TJ, Huerta P, Blum KI, Wilson MA (1996) Hippocampal CA1-region-restricted knockout of NMDAR1 gene disrupts synaptic plasticity, place fields, and spatial learning. Cold Spring Harb Symp Quant Biol 61:225–238.

Tsodyks M (1999) Attractor neural network models of spatial maps in hippocampus. Hippocampus 9:481–489.

Tsodyks MV, Skaggs WE, Sejnowski TJ, McNaughton BL (1996) Population dynamics and theta rhythm phase precession of hippocampal place cell firing: a spiking neuron model. Hippocampus 6:271–280.

Valero M, Navas-Olive A, de la Prida LM, Buzsaki G (2022) Inhibitory conductance controls place field dynamics in the hippocampus. Cell Rep 40:111232.

van Dijk MT, Fenton AA (2018) On How the Dentate Gyrus Contributes to Memory Discrimination. Neuron 98:832–845.

Whittington JCR, Muller TH, Mark S, Chen G, Barry C, Burgess N, Behrens TEJ (2020) The Tolman-Eichenbaum Machine: Unifying Space and Relational Memory through Generalization in the Hippocampal Formation. Cell 183:1249–1263 e1223.

Wilson CD, Serrano GO, Koulakov AA, Rinberg D (2017) A primacy code for odor identity. Nat Commun 8:1477.

Wilson MA, Tonegawa S (1997) Synaptic plasticity, place cells and spatial memory: study with second generation knockouts. Trends Neurosci 20:102–106.

Zaki Y, Pennington ZT, Morales-Rodriguez D, Bacon ME, Ko B, Francisco TR, LaBanca AR, Sompolpong P, Dong Z, Lamsifer S, Chen HT, Carrillo Segura S, Christenson Wick Z, Silva AJ, Rajan K, van der Meer M, Fenton A, Shuman T, Cai DJ (2025) Offline ensemble co-reactivation links memories across days. Nature 637:145–155.

Zhang SJ, Ye J, Miao C, Tsao A, Cerniauskas I, Ledergerber D, Moser MB, Moser EI (2013) Optogenetic dissection of entorhinal-hippocampal functional connectivity. Science 340:1232627.

Zipser D (1985) A computational model of hippocampal place fields. Behav Neurosci 99:1006–1018.

Ziv Y, Burns LD, Cocker ED, Hamel EO, Ghosh KK, Kitch LJ, El Gamal A, Schnitzer MJ (2013) Long-term dynamics of CA1 hippocampal place codes. Nat Neurosci 16:264–266.

