## Supplemental Figures and Tables for "Effective computations for hippocampal place cell phenomena in sparse untrained random networks"

### SUPPLEMENTAL MATERIAL

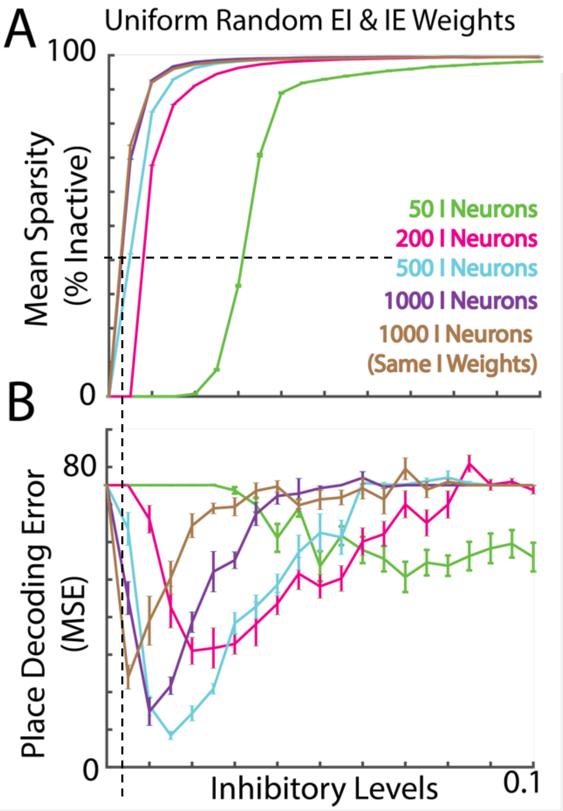


**Figure S1: Effect of I Population size and random inhibitory weights on sparsity and decoding error.** A) Network sparsity increases monotonically and saturates with high inhibition. E populations are fixed at 1000 E neurons. Results are averaged from ten simulations, each curve represents the percentage of active E neurons at time t, averaged over all time points in each simulation. Results using four different inhibitory (I) Neuron populations sizes. Also shown (brown) with equal E and I neurons (1000) but a single value for all inhibitory weights. Dotted lines reflect the critical inhibition (50% sparsity) for this configuration and its corresponding decoding error in (B). B) Decoding error is non-monotonic, with the lowest error occurring with 500 I neurons when EI and IE connections are random.

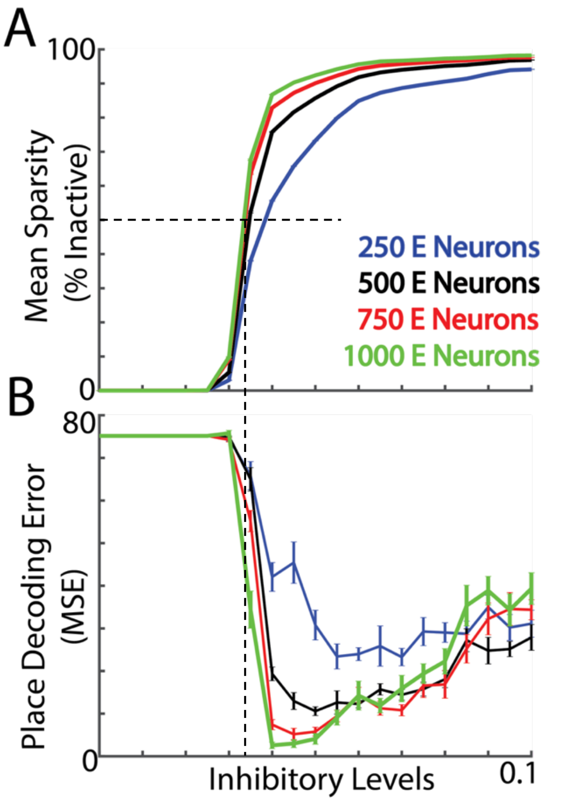


**Figure S2: Effect of E Population size on sparsity and decoding error.** A) Network sparsity increases monotonically and saturates with high inhibition. I populations are fixed at 50 I neurons with a single value for all EI and IE weights. Results are averaged from ten simulations, each curve represents the percentage of active E neurons at time t, averaged over all time points in each simulation. Results using four different excitatory (E) Neuron populations sizes are shown. Dotted lines reflect the critical inhibition (50% sparsity) for 1000 E neurons and the corresponding decoding error. B) Decoding error is non-monotonic, with a minimum at inhibition levels higher than the critical inhibition (50% Sparsity). Furthermore, the optimal inhibition level (where decoding error is minimized) approaches the critical inhibition as the E population increases.

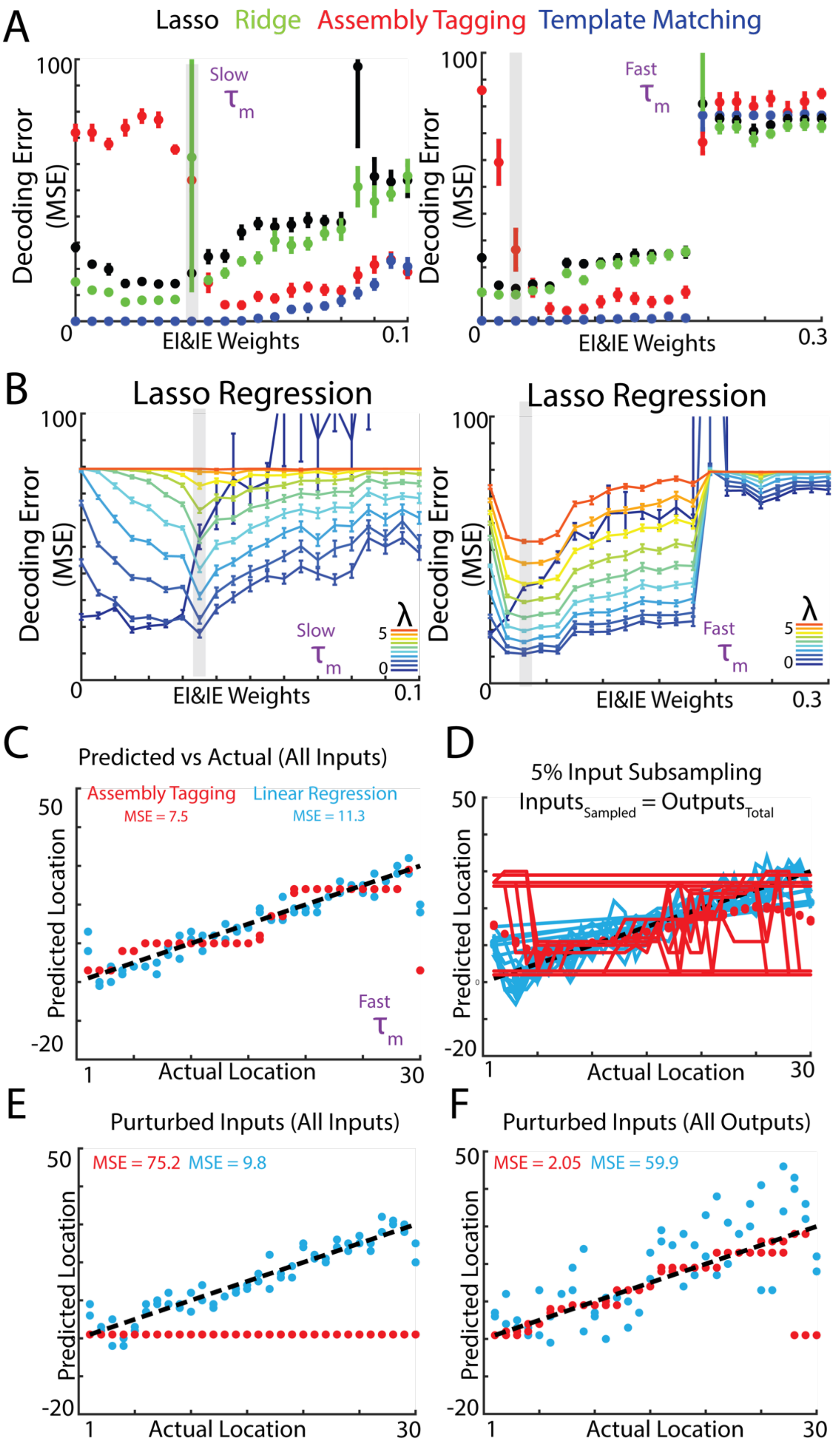

A
Lasso Ridge Assembly Tagging Template Matching
Decoding Error (MSE)
EI&IE Weights
Slow τm
Fast τm
B
Lasso Regression
λ
C
Predicted vs Actual (All Inputs)
Assembly Tagging
MSE = 7.5
Linear Regression
MSE = 11.3
Predicted Location
Actual Location
D
5% Input Subsampling
Inputs_Sampled = Outputs_Total
E
Purturbed Inputs (All Inputs)
MSE = 75.2 MSE = 9.8
F
Purturbed Inputs (All Outputs)
MSE = 2.05 MSE = 59.9

**Figure S3: Comparing multiple decoders.** A) Decoding error of four different decoders are shown (see methods). Note, the assembly tagging decoder is unique in that it performs poorly when network activity is saturated and performs well when activity is sparse. Left: LIF simulation with slow time constant parameters (see figure 2). Right: LIF simulation with fast time constant parameters. B) Effect of lasso regularization coefficient on the decoder performance using slow (left) or fast (right) LIF time constant parameters (See Fig 1H,I). Shown are results using regularization coefficient values between 0 and 5 ($\lambda$). C) Example decoding performance (actual vs decoded position) using the input activity prior to the LIF network layer. All input units (10,000) were used for decoding. Two decoders are being compared: Ordinary least-squared (OLS) linear regression (blue), and the assembly tagging decoder (red). D) Input subsampling degrades performance. When inputs are subsampled randomly to match the number of E neurons in the LIF network (500), there is increased variability in the decoding performance. Solid traces indicate individual examples from a random subsampling, and dots indicate the mean decoding performance across all examples. E) Same as (C), but with inputs perturbed by an additional phasic input. F) Example decoding performance using the LIF E neuron activity after perturbation of the inputs. Note that the assembly tagging decoder's performance is rescued at the expense of the linear decoder's performance.

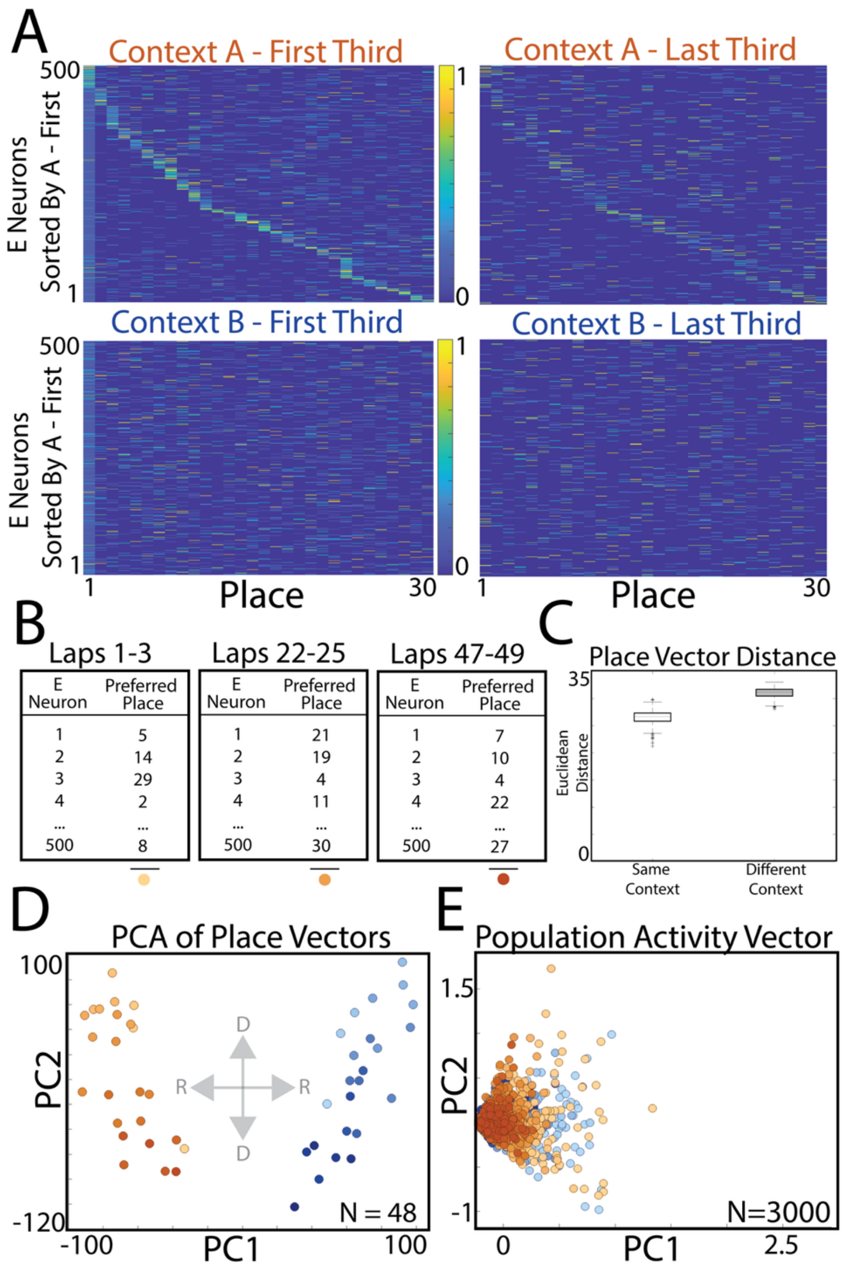


**Figure S4: Synaptic plasticity with a behavioral timescale (~1s analogous to behavioral time-scale plasticity (BTSP)) induces context-dependent drifting of the place representation**. A) Normalized rate maps across neurons for the first and last thirds of context A (top) and context B (bottom). Neurons are sorted according to the first third of the simulation in context A. B) Example of how place vectors can change over time due to the synaptic plasticity. C) Pairwise distance of all place vectors within the same context (A-A, B-B) and different context (A-B). Note that although both drift and remapping result in large pairwise distances in the high dimensional space (500 neurons), the former is smaller in comparison. D) 2D PCA of the place vectors from (C), colored by context and temporal order (light to dark). Note the disentangled directions of remapping (left and right (R)) vs drift (up and down (D)). E) The moment-to-moment population vector of activity fails to separate contexts, suggesting temporal coactivity is preserved even though place vectors are changing.

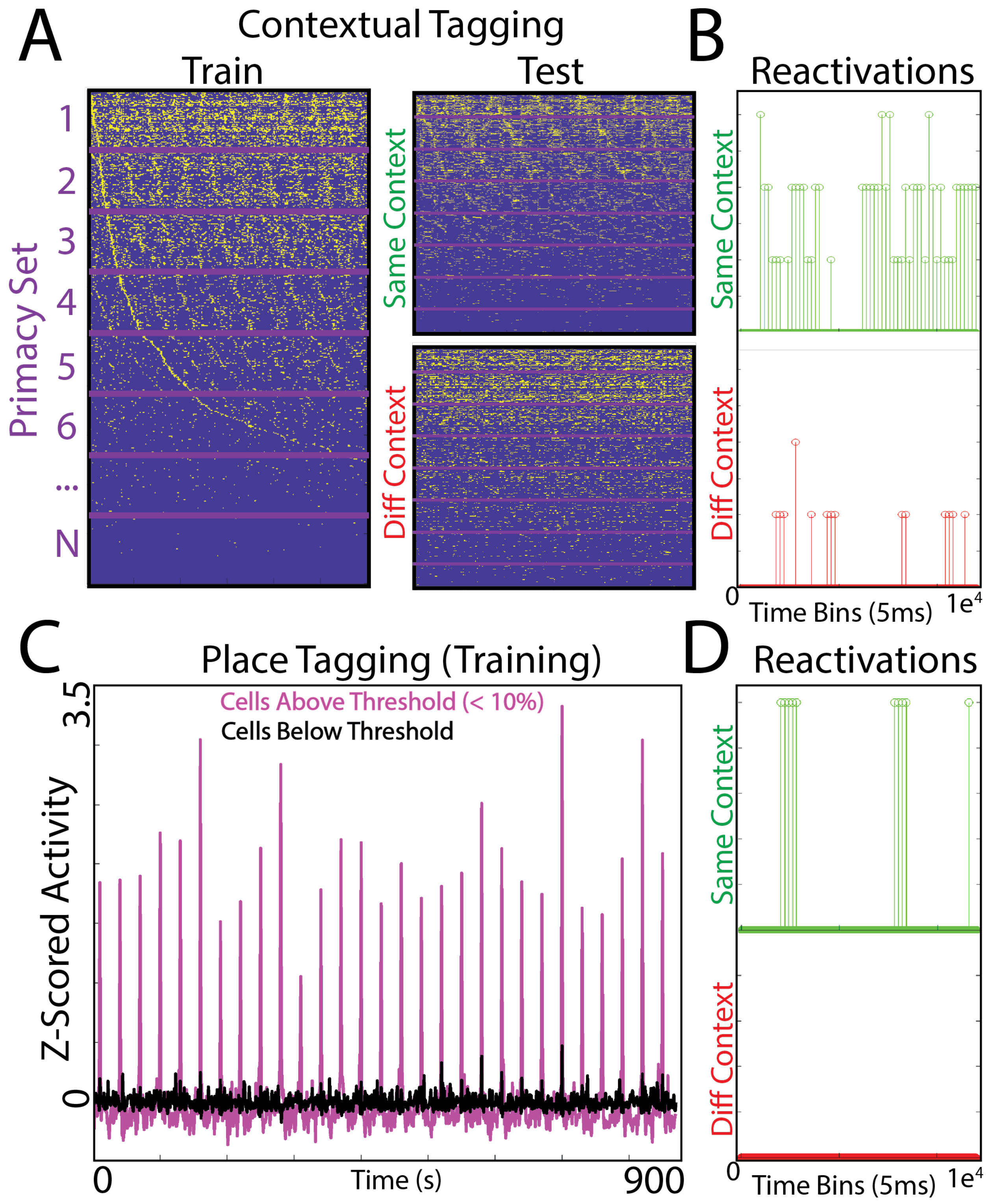

**Figure S5: Downstream tagging mechanism in LIF model for contextual and place signals.** A) In the Contextual Fear Conditioning task, the model is first exposed to the relevant context in order to retrieve the sorting of E neurons as they first reach 20 spikes (primacy sorting). From this sorting, neurons are grouped together for downstream readouts (Primacy Sets). Left: Spiking raster of E activity using primacy sorting of E neurons. Right: Spiking raster using the same sorting during a separate exposure to the same context or a different context. B) Downstream readouts detect when a primacy set is reactivated (>70% Neurons are active). Shown are the number of primacy sets activated at a given time point. Downstream motor responses can therefore generate "Freezing" when more than one primacy set is activated concurrently (Contextual Freezing). C) Place tagging mechanism: a special signal increases the activity of inputs at a special location, and these only affect a portion of the LIF E neurons. Only those neurons with activity crossing a threshold are tagged by downstream readouts. D) Reactivations of tagged E cell assembly in the same context vs a different context (Place Avoidance). See Fig. S6 for test simulations.

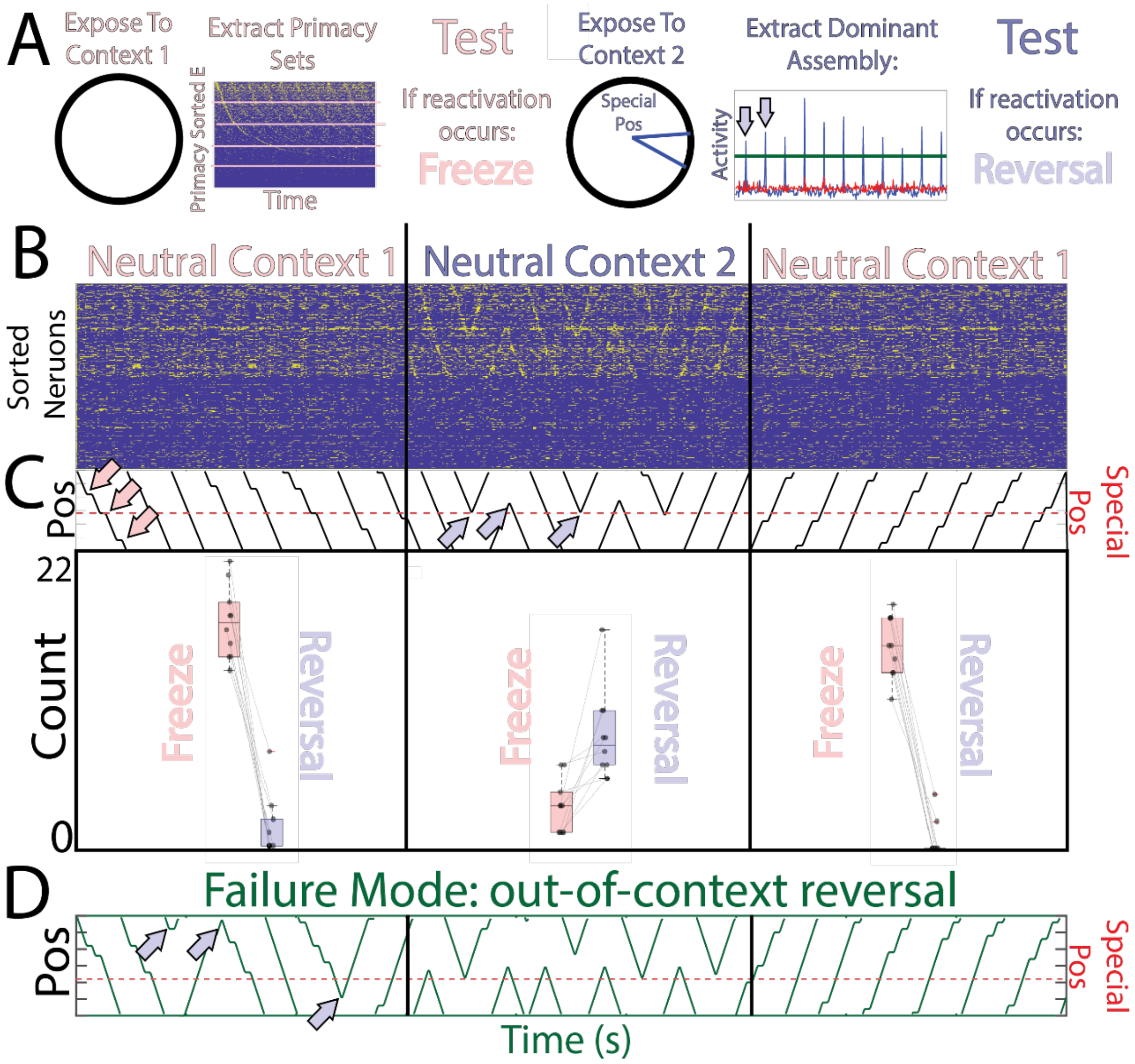


**Figure S6: Untrained EI network encodes contextual and positional information in sparse subpopulations tagged by downstream readouts** A) Left: Schematic of the encoding of special contexts: For any given input pattern, the network simulation will generate sequences of activity that can be used to generate primacy sets (see methods, Fig S5 A) grouping neurons based on spike timing. Right: Positional tagging: reinforced locations in a particular context are associated with increased E neuron activity in a subset of cells (blue arrows vs red curve) (see methods, Fig S5 C) . B) Top: Simulation of two neutral versions of the learned contexts during recall (no special context/location signals). Network output spikes are shown with neurons sorted according to positional selectivity in the avoidance context, note the remapping of location-specific tuning between the freeze and the avoid contexts. C)  Top: Position trajectories across the simulation, also shown are freeze (red arrows) and reversal (blue arrows) moments. Bottom: Scores of freeze and avoidance responses in the two contexts. E) Failure case of tagging results in memory-linking, whereby experience in one context is recollected in a different context.

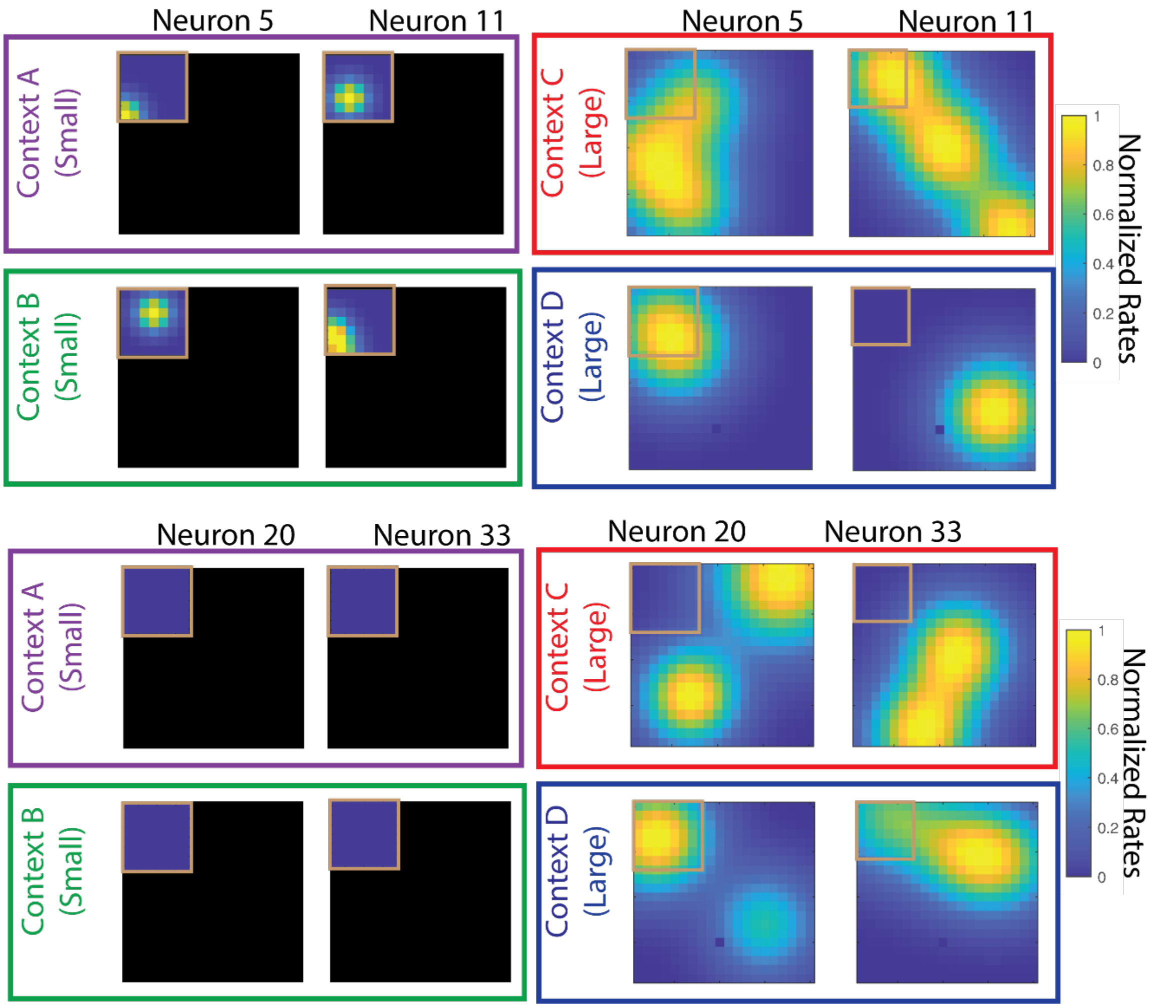


**Figure S7: Example place fields in small (Context A and B) and large (Context C and D) environments across contexts (Remapping).**

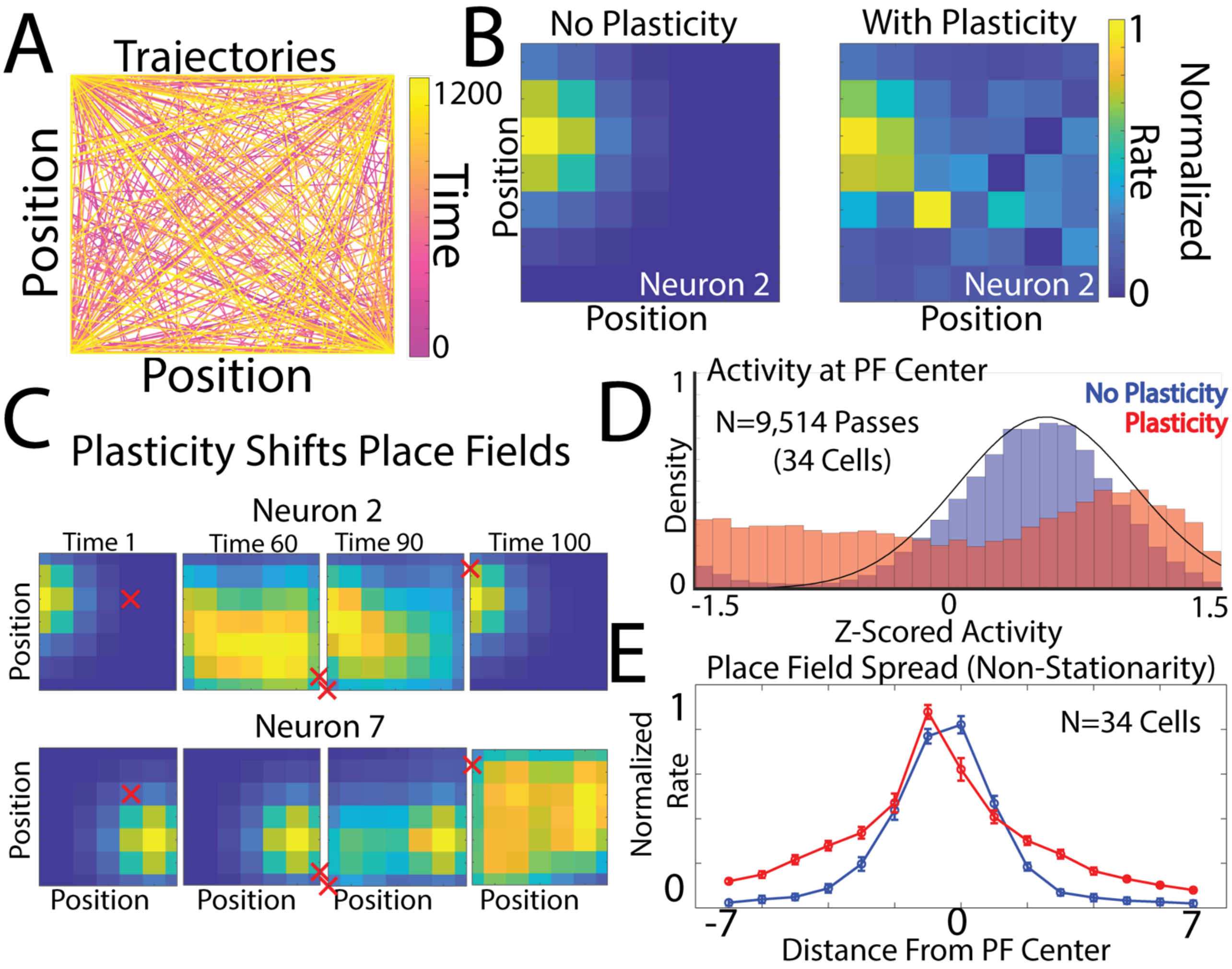


**Figure S8: Overdispersion examples and statistics.** A) Simulated trajectories in the 2D model, color coded according to the simulation time. The activity that occurs because of the trajectories produce changes in the network EE recurrent via a spike timing plasticity rule (STDP). B) Trial averaged rate maps of the same cell with and without recurrent plasticity in the network. C) Instantaneous rate maps of two cells, across 4 distinct time points, and the current location in the trajectory (red "x"). The peak activity is often near the trial averaged peak, but can shift as a result of changes in the recurrent weights, leading to activity that is over-dispersed from the center of the place field due to momentary non-local activity. D) Z-scored activity of multiple passes through a place field center with or without plasticity. The activity at the place field center is unreliable with plasticity. E) Trial-averaged activity is more spread across the space when plasticity rules are engaged.

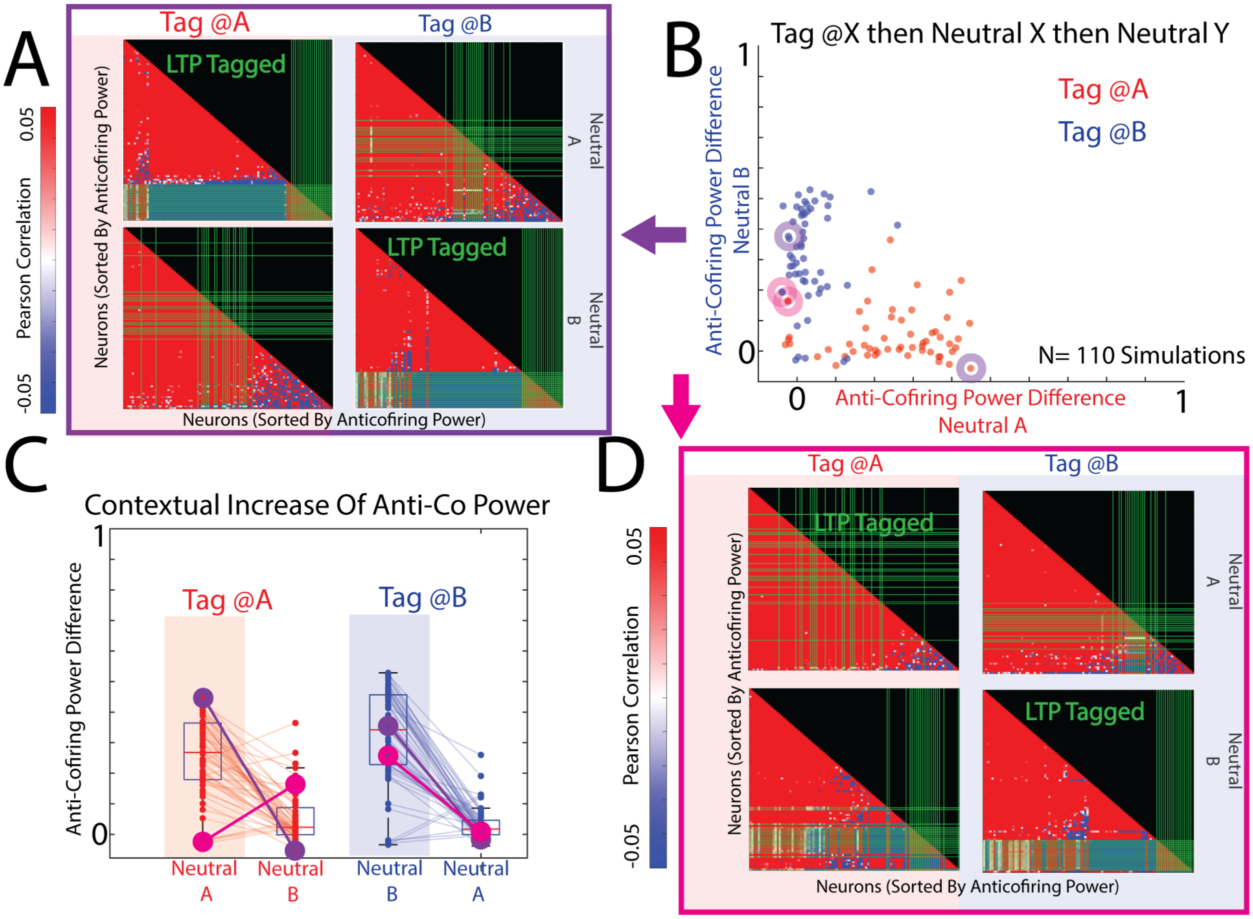


**Figure S9: Context-Specific Anti-Cofiring Statistics in the 2D DivSparse model:** A) Pearson correlation of network units after tagging and reexposure to the same (but neutral) context, compared to exposure to a novel neutral context. Green lines indicate the LTP tagged cells, which are more anti-cofiring in the respective tagged context. B) Scatterplot of the difference in mean anti-cofiring power between non-tagged vs tagged cells that were tagged in context A (red) or B (blue). After tagging, a neutral version of the same context was presented prior to remapping to a novel neutral context. Purple circles show the examples presented in (A), while magenta circles show outlier cases (D) that deviate from the expected trend of context-selective anti-cofiring. C) Boxplots of mean anti-cofiring difference in neutral context A or B after tagging in context A or B. D) Outlier case where one of the contexts does not generate context-selective anti-cofiring in the tagged context and another generates context-selective anti-cofiring in both contexts.

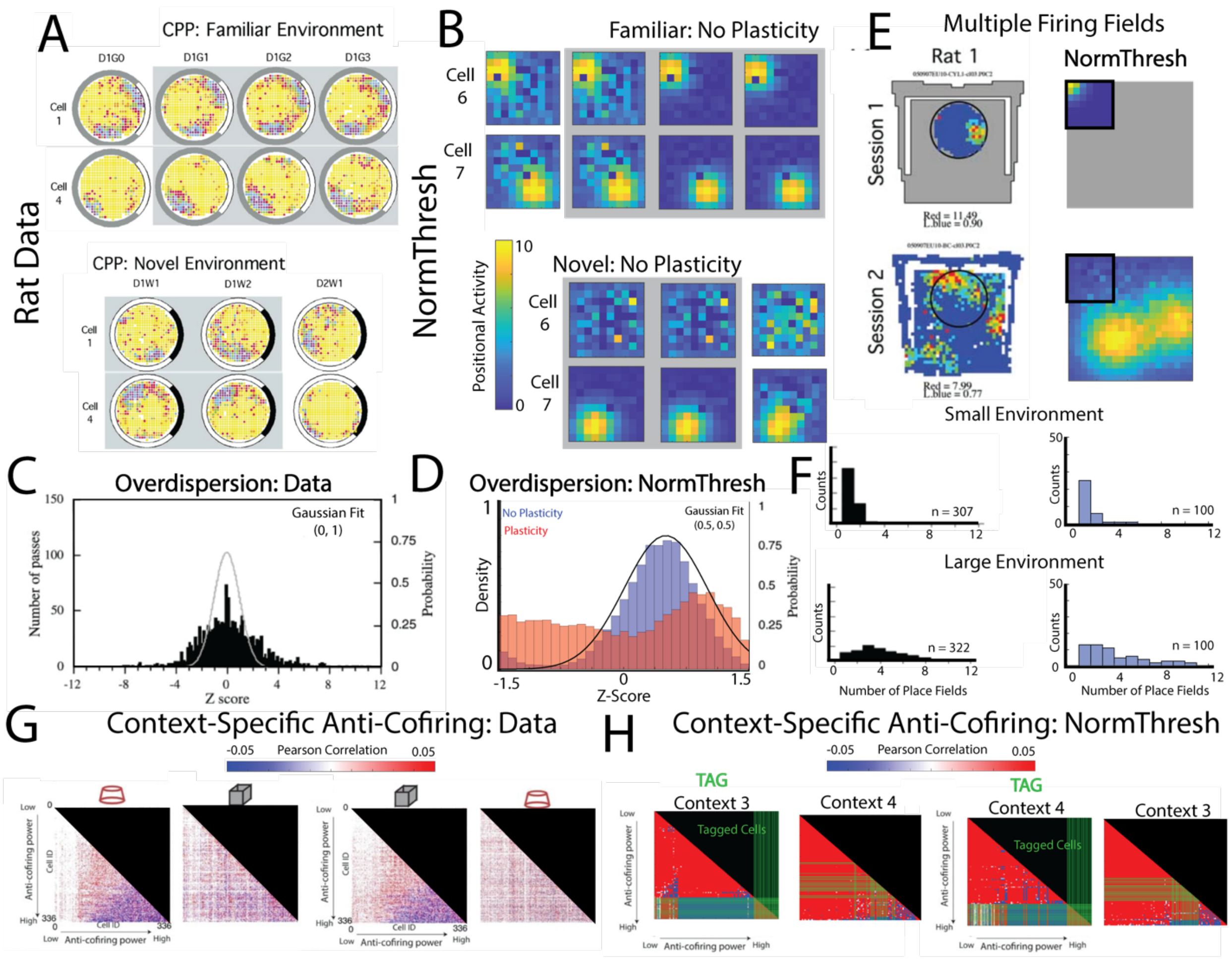


**Figure S10: DivSparse model reproduces several empirical HPC phenomena.** A) Adapted from Kentros et al. 1998, showing two cell's place fields in a familiar and novel environment after addition of NMDAR antagonist CPP (Gray Region). Place field formation and remapping is unaffected by CPP, although place field stability is sometimes altered by CPP in the novel environment. B) Analogous experiment as (A) but using the DivSparse simplified model with plasticity turned off (Gray Regions). Similar to the rat data, place field formation and remapping are unaffected by turning off plasticity, although place field stability also remains unaffected in the novel environment, marking a deviation in the model from the experimental data. C) Adapted from (Fenton and Muller, 1998), shown are z-scores of place cell activities as the animal makes momentary "passes" through the place field center. Also shown is a theoretical Gaussian function to reflect the overdispersion of activities across distinct passes through a place field. D) Analogous analysis as in (C) except using the DivSparse model with and without plasticity. Without plasticity, the Z-score of activities through multiple passes resembles a Gaussian distribution with reliable activity occurring with some variance resulting from noise added in the simulation. When plasticity is turned on, the same inputs lead to over dispersed activity that is not reliable across multiple passes through the place field. E) Adapted from (Fenton et al., 2008); Place field maps for the same cell in the experimental data (Left) or the DivSparse model (Right) showing the increased number of place fields when the environment is enlarged. Shown also are the relative sizes of the two environments with respect to each other. F) Histogram of the number of fields that result from the small (Top) and large environment (Bottom) for the data and the DivSparse model. G) Adapted from (Levy et al., 2023); Shown are the pairwise Kendall correlations for cells sorted according to the proportion of anti-cofiring synaptic partners (correlation < -0.05). Note that the anti-cofiring distribution is context specific, leading to a scrambling of the pairwise sorting across contexts (Circle vs Square environment). H) The analogous analysis is shown for the DivSparse model, where experience dependent LTP was induced (as in Fig. 4 and Fig S9) to demonstrate the context specificity that arises in the anti-cofiring pairwise distributions. Note also that the cells that end up tagged by LTP (green lines) are more likely to be cells with high anti-cofiring power.

### REFERENCES

Fenton AA, Muller RU (1998) Place cell discharge is extremely variable during individual passes of the rat through the firing field. Proc Natl Acad Sci U S A 95:3182-3187.

Fenton AA, Kao H-Y, Neymotin SA, Olypher AV, Vayntrub Y, Lytton WW, Ludvig N (2008) Unmasking the CA1 ensemble place code by exposures to small and large environments: more place cells and multiple, irregularly-arranged, and expanded place fields in the larger space. J Neurosci 28:11250-11262.

Levy ERJ, Carrillo-Segura S, Park EH, Redman WT, Hurtado JR, Chung S, Fenton AA (2023) A manifold neural population code for space in hippocampal coactivity dynamics independent of place fields. Cell Reports 42.
